# YAP activity is necessary and sufficient for basal progenitor abundance and proliferation in the developing neocortex

**DOI:** 10.1101/416537

**Authors:** Milos Kostic, Judith T.M.L. Paridaen, Katherine Long, Nereo Kalebic, Barbara Langen, Pauline Wimberger, Hiroshi Kawasaki, Takashi Namba, Wieland B. Huttner

**Affiliations:** Max Planck Institute of Molecular Cell Biology and Genetics, Pfotenhauerstrasse 108, 01307 Dresden, Germany; Technische Universität Dresden, Universitätsklinikum Carl Gustav Carus, Klinik und Poliklinik für Frauenheilkunde und Geburtshilfe, Fetscherstraße 74, D-01307 Dresden, Germany.; Department of Medical Neuroscience, Graduate School of Medical Sciences, Kanazawa University, Ishikawa 920-8640, Japan

**Keywords:** Hippo signaling, YAP activity, neocortical evolution, SVZ, basal progenitors

## Abstract

The expansion of the neocortex during mammalian evolution has been linked to an enlargement of the subventricular zone during cortical development and an increase in the proliferation of the basal progenitors residing therein. Here, we explored a potential role of YAP, the major downstream effector of the Hippo signaling pathway, in proliferation of basal progenitors. We show that YAP expression and activity are high in ferret and human basal progenitors, which are known to exhibit high proliferative capacity, but low in mouse basal progenitors, which lack such capacity. To induce YAP activity in mouse basal progenitors, we expressed a constitutively active YAP (CA-YAP). This resulted in an increase in proliferation of basal progenitor. In addition, CA-YAP expressing mouse basal progenitors promoted the production of upper-layer neurons. To investigate if YAP is required for the proliferation of basal progenitors, we pharmacologically interfered with the function of YAP in the developing ferret and human neocortex. This resulted in a decrease of cycling basal progenitors. In concert, genetical interference with the function of YAP in ferret developing neocortex resulted in decreased abundance of basal progenitors. Together, our data indicate that YAP promotes the proliferation of basal progenitors and suggest that changes in YAP activity levels contributed to the evolutionary expansion of the neocortex.

## Introduction

The neocortex, the seat of higher cognitive functions, undergoes substantial expansion during the evolution of certain mammalian brains such as human. A major factor in neocortical expansion, notably regarding the increase in the number of cortical neurons, is thought to be an increased proliferative capacity of cortical neural progenitor cells (cNPCs) (Fietz and Huttner, 2011; Florio and Huttner, 2014; Geschwind and Rakic, 2013; Lui et al., 2011; Namba and Huttner, 2017; Rakic, 2009; Wilsch-Bräuninger et al., 2016).

Two principal classes of cNPCs exist in the developing neocortex, referred to as apical progenitors (APs) and basal progenitors (BPs) (Florio and Huttner, 2014; Lui et al., 2011; Namba and Huttner, 2017). The defining feature of APs is that they undergo mitosis at the ventricular (apical) surface of the ventricular zone (VZ), the primary germinal zone where the AP cell bodies reside (Florio and Huttner, 2014; Namba and Huttner, 2017). At the onset of neurogenesis, the major AP cell type are apical (or ventricular) radial glia (aRG) (Fietz and Huttner, 2011; Florio and Huttner, 2014; Lui et al., 2011; Namba and Huttner, 2017). The defining feature of BPs is that they undergo mitosis away from the apical surface, typically in a secondary germinal zone called the subventricular zone (SVZ) where the BP cell bodies reside (Haubensak et al., 2004; Miyata et al., 2004; Noctor et al., 2004). BPs originate from APs, delaminate from the apical surface, migrate beyond the VZ, and thus form the SVZ. There are two main types of BPs, basal intermediate progenitor (bIPs) and basal (or outer) radial glia (bRG). In contrast to aRG, which are epithelial cells exhibiting apical-basal polarity with contact to the ventricle and (in the canonical form) to the basal lamina, bIPs are nonepithelial cells that no longer exhibit apical-basal polarity and that have lost contact to both the ventricle and the basal lamina (Haubensak et al., 2004; Miyata et al., 2004; Noctor et al., 2004). bRG, however, though lacking an apical process that reaches the ventricle, retain epithelial features in that they (in the canonical form) possess a basal process that contacts the basal lamina (Fietz et al., 2010; Hansen et al., 2010; Reillo et al., 2011).

BP composition and proliferative capacity may differ greatly between a developing lissencephalic neocortex (e.g. mouse) and a developing gyrencephalic neocortex (e.g. ferret and human). In the developing mouse neocortex BPs mostly comprise bIPs that typically undergo neurogenic consumptive divisions giving rise to two neurons; compared to the aRG they derive from, these neurogenic bIPs characteristically upregulate the transcription factor Tbr2 and downregulate the transcription factor Sox2 (Haubensak et al., 2004; Miyata et al., 2004; Noctor et al., 2004). Only a minor portion of mouse BPs are bRG, and their proliferative potential is limited (Shitamukai et al., 2011; Wang et al., 2011). In contrast, in the developing ferret and human neocortex, the majority of BPs are proliferative bRG (highly proliferative in human) that do not express Tbr2 but rather maintain expression of Sox2 (Fietz et al., 2010; Hansen et al., 2010; Reillo et al., 2011). Moreover, as first shown in a seminal contribution for the developing monkey neocortex (Smart et al., 2002), the SVZ in a gyrencephalic neocortex is characteristically split into two morphologically distinct zones, an inner SVZ (iSVZ) and an outer SVZ (oSVZ). Of note, the evolutionary expansion of the neocortex has been linked to an increase in the proliferative capacity and abundance of BPs in the oSVZ, especially of bRG (Dehay et al., 2015; Fietz and Huttner, 2011; Lui et al., 2011; Namba and Huttner, 2017). However, the molecular players that differentially promote the proliferative capacity of BPs across the various mammalian species remain largely unknown.

To gain insight into this issue, we examined a major molecular mechanism known to regulate organ size, the Hippo/YAP signaling pathway (Barry and Camargo, 2013; Camargo et al., 2007; Lian et al., 2010; Yu et al., 2015). The core of Hippo/YAP signaling is the YAP protein, whose transcriptional activity is regulated by phosphorylation. Phosphorylated YAP (phospho-YAP) is retained in the cytoplasm, whereas dephosphorylated YAP (active YAP) translocates to the nucleus and activates the expression of genes linked to cell proliferation (Zanconato et al., 2015; Zhao et al., 2010; Zhao et al., 2007; Zhao et al., 2008). Recent studies dissecting the roles of the cadherin family members Dchs1 and Fat4 (Cappello et al., 2013) and of the tumor suppressor neurofibromatosis 2 (Lavado et al., 2013; Lavado et al., 2014) and investigating heterotopia formation (Saito et al., 2017) in mouse brain development have reported that YAP promotes the proliferation of mouse APs. These studies, however, have not focused on a potential role of YAP in regulating the proliferation of BPs, nor have they addressed the question whether differences in YAP activity may underlie the differences in the proliferative capacity of BPs across various mammalian species in the context of the evolutionary expansion of the neocortex.

In the present study, we have identified differences in YAP expression and YAP activity between the developing lissencephalic mouse and gyrencephalic ferret and human neocortex that match the differences in the proliferative capacity of BPs across these species. Enhancing YAP activity in mouse BPs induced their proliferation and therefore shifted their fate from neurogenic to proliferative. In contrast, inhibition of endogenous YAP activity by verteporfin or by administration of a dominant-negative YAP reduced BP proliferation in developing ferret and human neocortex. Taken together, these findings suggest that an upregulation of YAP activity contributed to the increased proliferative capacity of BPs in the context of the evolutionary expansion of the neocortex.

## Results

### BPs with high proliferative capacity, which are abundant in embryonic ferret and fetal human neocortex but are lacking in embryonic mouse neocortex, show high YAP expression

To explore whether YAP is differentially expressed between the principal germinal zones of developing mouse and human neocortex, the VZ vs. the SVZ, and between the major classes of cNPCs therein, the APs vs. BPs, respectively, we first analyzed the levels of *Yap*/*YAP* mRNA in these germinal zones and cNPC classes that have been determined in previously published transcriptome analyses (Figure S1) (Fietz et al., 2012; Florio et al., 2015). *Yap*/*YAP* mRNA was robustly expressed in the VZ of both embryonic day (E) 14.5 mouse and 13 week-post-conception (wpc) human neocortex (Figure S1A), and accordingly in mouse and human aRG (Figure S1B), the major AP type (Namba and Huttner, 2017). Strikingly, *Yap*/*YAP* mRNA was found to be expressed in the human iSVZ and oSVZ, but not the mouse SVZ (Figure S1A), and in human bRG, but not mouse BPs (Figure S1B). Given that both human and mouse APs, and human, but not mouse, BPs are endowed with the ability to expand their population size by cell proliferation (Namba and Huttner, 2017), these data provided a first indication that the proliferative capacity of cNPCs, notably of BPs, may be linked to the expression of YAP. Consistent with this notion, no significant *Yap*/*YAP* mRNA expression was detected in the mouse and human cortical plate (CP) (Figure S1A), a zone of the developing neocortex lacking cell proliferation, nor in post-mitotic neurons (Figure S1B; note that the *YAP* mRNA in the human neuron fraction reflects the presence of bRG in G1 in this fraction (Florio et al., 2015)).

Further evidence in support of a potential role of YAP in cNPC proliferation was provided by the comparison of *Yap* mRNA levels between *Tis21*-GFP–negative and *Tis21*-GFP–positive aRG of E14.5 mouse neocortex, determined previously (Florio et al., 2015) using the *Tis21-*GFP knock-in (*Tis21^tm2(Gfp)Wbh^*) mouse line (Haubensak et al., 2004). This revealed a two-fold higher *Yap* mRNA level in *Tis21*-GFP–negative aRG as compared to *Tis21*-GFP–positive aRG (Figure S1C) (Florio et al., 2015), in line with both cNPC types being capable of self-renewal but only the former cNPC subpopulation being capable of expansion by proliferation (Attardo et al., 2008; Haubensak et al., 2004). Finally, comparison of mRNA levels between a prospective gyrus vs. a prospective sulcus of developing (postnatal day (P) 2) ferret neocortex, available in a previously published transcriptome dataset (De Juan Romero et al., 2015), showed that the *Yap* mRNA level was higher in the oSVZ of the prospective gyrus than the prospective sulcus (Figure S1D), consistent with the notion that a relative increase in cNPC proliferation in this germinal zone contributes to gyrus formation (Hansen et al., 2010; Reillo et al., 2011; Wang et al., 2011). Taken together, these *Yap*/*YAP* mRNA data raised the possibility that YAP may not only have a role in the proliferation of APs, as previously shown for embryonic mouse neocortex (Lavado et al., 2013; Lavado et al., 2014), but that differences in the level of active YAP may underlie the differences in the proliferative capacity of mouse vs. ferret and human BPs.

We therefore examined the expression of the YAP protein in embryonic mouse, embryonic ferret and fetal human neocortex by immunofluorescence (Figure 1A-C, F-H). Consistent with the mRNA expression data (Figure S1A), YAP immunoreactivity was overt in the E14.5 mouse, E36 ferret and 14 wpc human VZ and in the ferret and human SVZ, notably the oSVZ, but was low in the mouse SVZ (Figure 1A-C). In the case of the embryonic ferret oSVZ, the YAP immunostaining revealed cells exhibiting a basal process (Figure 1B’), suggesting that they were bRG.

**Figure 1.**
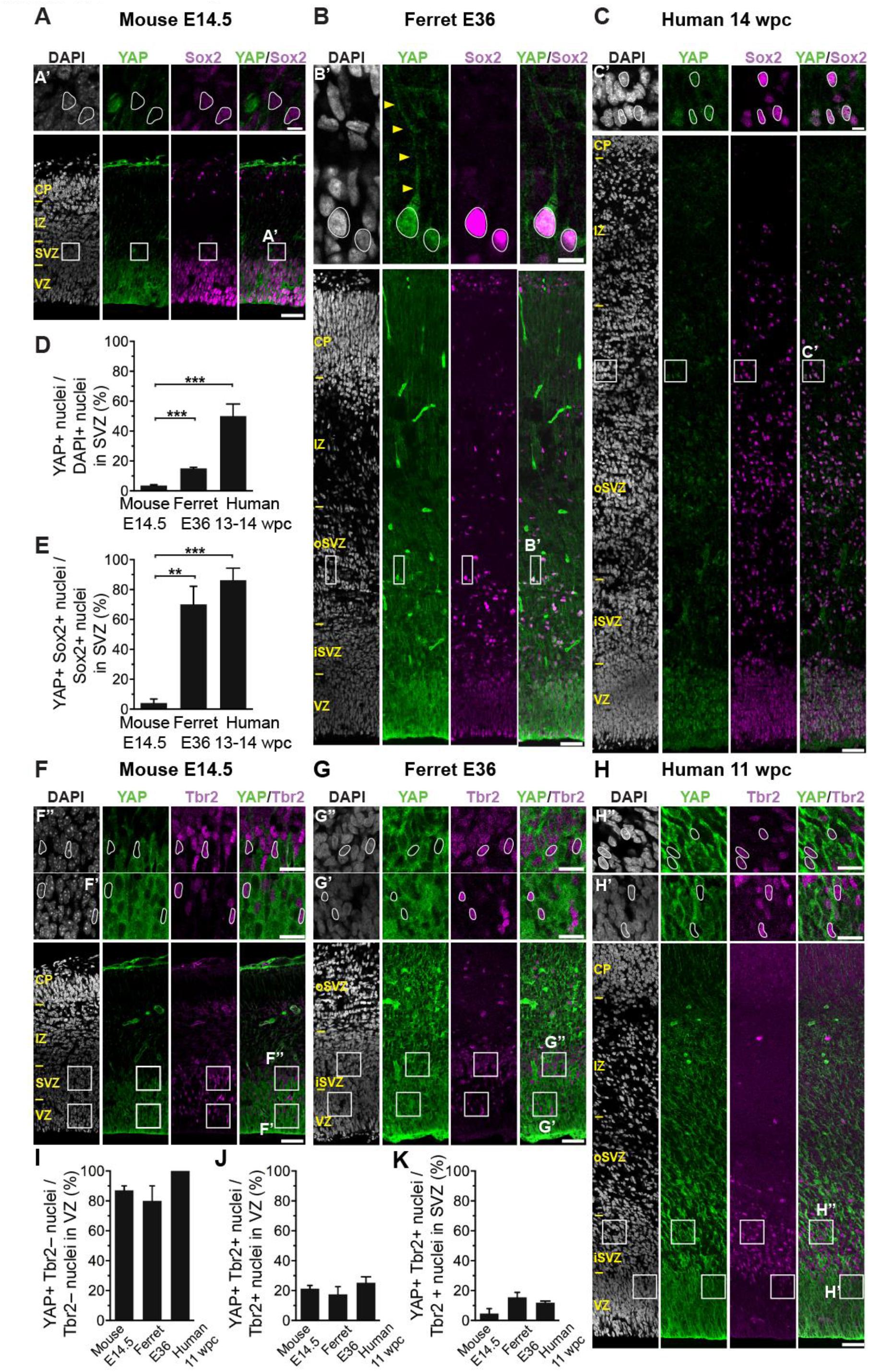
The majority of ferret and human, but not mouse, Sox2-positive Tbr2-negative BPs exhibit nuclear YAP. See also Figures S1, S2. (A-C) Double immunofluorescence for YAP (green) and Sox2 (magenta), combined with DAPI staining (white), of mouse E14.5 (A), ferret E36 (B) and human 14 wpc (C) neocortex. Boxes indicate areas in the SVZ (A) and oSVZ (B, C) that are shown at higher magnification in panels A’, B’ and C’; selected Sox2-positive nuclei that are YAP-negative in mouse and YAP-positive in ferret and human are outlined by white lines; arrowheads indicate a YAP-positive basal process of a bRG. (D, E) Quantification of the percentage of DAPI-stained nuclei (D) and Sox2-positive nuclei (E) in the SVZ that are YAP-positive in mouse E14.5, ferret E36 and human 13-14 wpc neocortex. Two-three images per embryo/fetus were taken, 30 randomly picked DAPI-stained nuclei (D) and Sox2-positive nuclei (E) in the SVZ were scored per image, and the values obtained were averaged for each embryo/fetus. Data are the mean of 4 embryos/fetuses. (F-H) Double immunofluorescence for YAP (green) and Tbr2 (magenta), combined with DAPI staining (white), of mouse E14.5 (F), ferret E36 (G) and human 11 wpc (H) neocortex. Boxes indicate areas in the VZ and SVZ (F) or iSVZ (G, H) that are shown at higher magnification in panels F’, F”, G’, G’’, H’ and H’’, as indicated; selected Tbr2-positive nuclei that are YAP-negative in mouse, ferret and human are outlined by white lines. (I-K) Quantification of the percentage of Tbr2-negative nuclei in the VZ (I), Tbr2-positive nuclei in the VZ (J) and Tbr2-positive nuclei in the SVZ (K) that are YAP-positive in mouse E14.5, ferret E36 and human 11 wpc neocortex. Two-three images per embryo/fetus were taken, 30 randomly picked Tbr2-negative nuclei in the VZ (I) and Tbr2-positive nuclei in the VZ (J) and SVZ (K) were scored per image, and the values obtained were averaged for each embryo/fetus. Data are the mean of 3-4 embryos/fetuses. (A-C, F-H) Images are 1-µm optical sections. Scale bars, 50 µm in (A-C, F-H), 10 µm in (A’, B’, C’), 20 µm in (F’, F”, G’, G’’, H’, H’’). (D, E, I-K) Error bars indicate SEM; ** *P* <0.01, *** *P* <0.001 (one-way ANOVA, post-hoc Tukey HSD).

### YAP-positive BPs in embryonic ferret and fetal human neocortex express Sox2 rather than Tbr2, and their YAP is nuclear and active

For YAP to be able to promote the expression of genes linked to proliferation, it needs to be nuclear and unphosphorylated (Zanconato et al., 2015; Zhao et al., 2010; Zhao et al., 2007; Zhao et al., 2008). Interestingly, in line with a potential role of YAP in BP proliferation, the 13-14 wpc human SVZ contained the highest percentage of YAP-positive nuclei (50%), followed by the E36 ferret SVZ (15%), whereas the E14.5 mouse SVZ exhibited only low levels of YAP-positive nuclei (4%) (Figure 1D).

We sought to obtain direct evidence that the higher level of nuclear YAP protein seen in embryonic ferret and fetal human SVZ as compared to mouse SVZ (Figure 1D) is linked to the increased proliferative capacity of ferret and human BPs vs. mouse BPs. To this end, we compared the expression of YAP with that of Sox2 (Figure 1A-C), an indicator of proliferative capacity (Hansen et al., 2010; Reillo et al., 2011; Wang et al., 2011). Specifically, we asked whether SVZ nuclei positive for Sox2 are also positive for YAP. Indeed, the vast majority of the Sox2-positive BP nuclei in the E36 ferret and 13-14 wpc human SVZ were YAP-positive (70% and 86%, respectively), whereas this was the case for only a minor proportion of the Sox2-positive nuclei in the E14.5 mouse SVZ (4%) (Figure 1E).

We next investigated whether the YAP protein in the Sox2-positive BP nuclei was in an active form, i.e. in a dephosphorylated state. A first, qualitative indication that this was the case was obtained by comparing the immunofluorescence signals for total YAP and phospho-YAP in Sox2-positive BP nuclei in the SVZ of human 11 wpc neocortex (Figure S2A). Relative to the cytoplasmic signals for total YAP and phospho-YAP, respectively, this comparison showed a much lower signal for nuclear phospho-YAP than nuclear total YAP, suggesting that most nuclear YAP in fetal human neocortical BPs was in the dephosphorylated state and hence active. We therefore used this approach to compare the levels of active YAP in Sox2-positive BP nuclei in the SVZ of mouse E13.5, ferret E36 and human 11 wpc neocortex. To this end, we subtracted the internal standard-adjusted immunofluorescence signal for nuclear phospho-YAP from that of nuclear total YAP to obtain information about nuclear dephospho-YAP levels (see legend to Figure S2B for details). This revealed substantially higher levels of active YAP in the Sox2-positive BP nuclei of ferret E36 and human 11 wpc neocortex than mouse E13.5 neocortex (Figure S2B).

We complemented these data by determining the effect of lambda protein phosphatase treatment of cryosections on the YAP immunofluorescence signals in Sox2-positive BP nuclei in the SVZ of mouse E14.5, ferret E36 and human 12-13 wpc neocortex. For YAP immunofluorescence, two rabbit monoclonal antibodies were used together, one recognizing YAP irrespective of serine127 phosphorylation (total YAP) and the other recognizing the serine127 phosphorylation site when phosphorylated (phospho-YAP). Hence, in the control, i.e. without lambda protein phosphatase treatment, the YAP immunofluorescence signal reflects the binding of primary antibodies to 1-2 sites, depending on whether serine127 is phosphorylated or not. In the case of mouse, lambda protein phosphatase treatment reduced the YAP immunofluorescence signal by ≈40% compared to control (Figure S2C). As lambda protein phosphatase treatment resulted in complete dephosphorylation of serine127 (see Methods), this ≈40% reduction suggested that about two-thirds (40/60) of the YAP protein in the Sox2-positive BP nuclei of mouse E14.5 neocortex was in phosphorylated form, and thus inactive. In contrast, in the case of ferret and human, lambda protein phosphatase treatment reduced the YAP immunofluorescence signal by only ≈20% and 10%, respectively, compared to the controls (Figure S2C), consistent with only ≈25% (20/80) and ≈10% (10/90) of the YAP protein in the Sox2-positive BP nuclei of ferret E36 neocortex and human 12-13 wpc neocortex, respectively, being in phosphorylated form. These data therefore corroborated our finding that there are markedly higher levels of active YAP in the Sox2-positive BP nuclei of embryonic ferret and fetal human than embryonic mouse neocortex.

Results essentially opposite to the occurrence of YAP in Sox2-positive BP nuclei were obtained when we compared the expression of YAP with that of Tbr2 (Figure 1F-H), which in embryonic mouse neocortex has been established as a marker of differentiating BPs, notably of neurogenic bIPs (Englund et al., 2005; Kowalczyk et al., 2009; Pontious et al., 2008). Thus, only a minor proportion (<16%) of the Tbr2-positive nuclei in the E14.5 mouse, E36 ferret and 11 wpc human SVZ were also positive for YAP (Figure 1K). Similarly, only ≈20% of the Tbr2-positive nuclei in the VZ of developing mouse, ferret and human neocortex, which constitute newborn BPs (Arai et al., 2011), were YAP-positive (Figure 1J). In contrast, the overwhelming majority (>75%) of the Tbr2-negative nuclei in the VZ of developing mouse, ferret and human neocortex, which constitute APs capable of expansion by proliferation, were YAP-positive (Figure 1I).

Taken together, these data suggest that the proliferative capacity of cNPCs, notably of BPs, correlates with the occurrence of nuclear YAP activity, with a high percentage of proliferating APs in developing mouse, ferret and human neocortex and of proliferating BPs in embryonic ferret and fetal human neocortex exhibiting nuclear (and mostly active) YAP, whereas this was the case for only a low percentage of the proliferating BPs in embryonic mouse neocortex. Conversely, only a low percentage of the nuclei of differentiating (rather than proliferating) BPs in the developing mouse, ferret and human neocortex exhibited nuclear YAP (and hence nuclear YAP activity, if any).

### Conditional expression of constitutively active YAP in embryonic mouse neocortex decreases Tbr2-positive BPs and increases Sox2-positive BPs

To examine if YAP activity is functionally linked to the proliferative capacity of BPs, we conditionally expressed a constitutively active YAP (CA-YAP) in BPs of the embryonic mouse neocortex, which lack proliferative potential and in which YAP expression normally is very low (Figure 1A,D,E). In the CA-YAP, two serine residues were replaced by alanine (S112A and S382A), mutations that have been shown to stabilize and to increase the nuclear localization of the YAP protein (Camargo et al., 2007; Dong et al., 2007; Zhao et al., 2010). The plasmid used for CA-YAP expression (referred to as CA-YAP–expressing plasmid) contained a strong constitutive promoter (CAGGS) driving a floxed membrane EGFP followed by the CA-YAP and an IRES-linked nuclear RFP reporter (Figure 2A left). Expression of CA-YAP and RFP upon Cre-mediated excision of the floxed EGFP was validated by transfection of HEK293T cells (Figure S3A) The same plasmid but lacking the CA-YAP module was used as control (Figure 2A left). Conditional expression of CA-YAP in mouse BPs was achieved by in utero electroporation (IUE) of the neocortex of E13.5 embryos of the *Tis21*-CreER^T2^ knock-in (*Btg2^tm1.1(cre/ERT2)Wbh^*) mouse line (Wong et al., 2015) (Figure 2A right). In embryos of this mouse line, the expression of tamoxifen-activated Cre follows that of Tis21, that is, is specific for BP-genic aRG and BPs (Wong et al., 2015). Conditional CA-YAP expression in mouse neocortical BPs by this approach was validated by analysis of E14.5 embryos (Figure S3B,C). Furthermore, conditional CA-YAP expression by this approach drove expression of a previously identified YAP target gene, CTGF (Malik et al., 2015; Zanconato et al., 2015; Zhao et al., 2008), in BPs three days after IUE (Figure S3B,D; for details see supplementary text).

**Figure 2.**
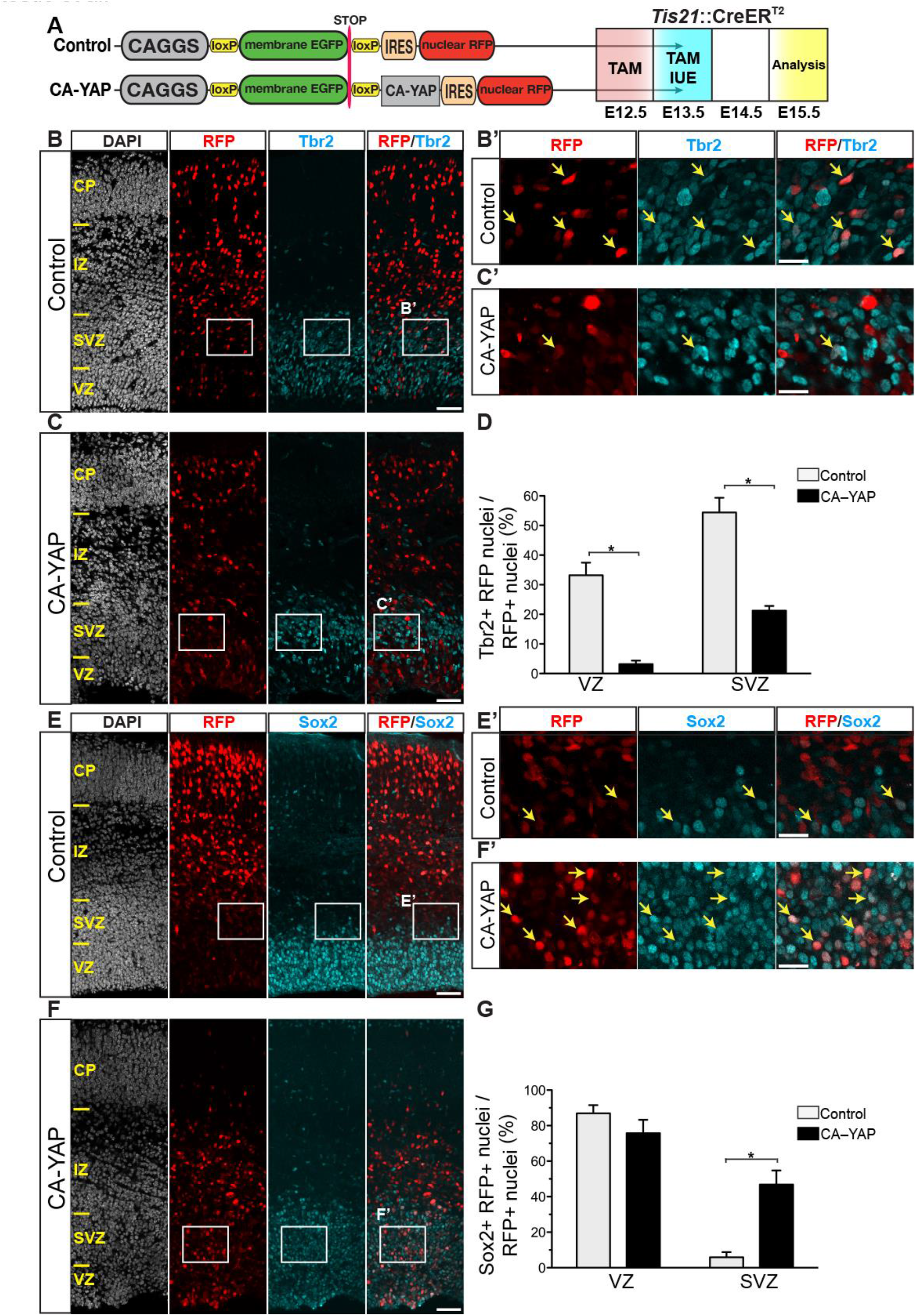
Conditional CA-YAP expression in the BP-genic lineage of embryonic mouse neocortex decreases production of Tbr2-positive BPs and increases generation of Sox2-positive BPs See also Figures S3 and S5. (A) Left, cartoon showing control (top) and CA-YAP–expressing (bottom) plasmid. Right, flow scheme of experiments. *Tis21*::CreER^T2^ heterozygous mouse embryos received tamoxifen (TAM) at E12.5 and E13.5, and the neocortex was *in utero* electroporated (IUE) at E13.5 with control plasmid (B, D, E, G) or CA-YAP–expressing plasmid (C, D, F, G) followed by analysis at E15.5. (B, C, E, F) Double immunofluorescence for RFP (red) and either Tbr2 (B, C) or Sox2 (E, F) (cyan), combined with DAPI staining (white). Boxes indicate areas in the SVZ that are shown at higher magnification in panels B’, C’, E’ and F’; arrows indicate selected RFP-positive nuclei that are Tbr2-positive (B’, C’) or Sox2-positive (E’, F’). Images are 1-µm optical sections. Scale bars, 50 µm in (B, C, E, F), 20 µm in (B’, C’, E’, F’). (D, G) Quantification of the percentage of RFP-positive nuclei that are Tbr2-positive (D) and Sox2-positive (G) in the VZ and SVZ, upon control (light grey) and CA-YAP (black) electroporation. Two images (1-µm optical sections), each of a 200 µm-wide field of cortical wall, per embryo were taken, and the percentage values obtained were averaged for each embryo. Data are the mean of 4 embryos from four separate litters. The mean ± SEM is shown; * *P* <0.01 (Mann-Whitney *U*-test).

We first examined the effects of conditional CA-YAP expression on BP fate, by analyzing Tbr2 and Sox2 expression 2 days after IUE of *Tis21*-CreER^T2^ mouse embryos (Figure 2A-G). This revealed, among the RFP-positive progeny of the targeted cells, a marked reduction in the proportion of Tbr2-positive cells in the VZ and SVZ (Figure 2D) and a striking increase in the proportion of Sox2-positive cells in the SVZ but not VZ (Figure 2G), compared to control. Hence, increasing YAP activity in neocortical BPs of embryonic mouse is sufficient to change their fate, interfering with their differentiation to a neurogenic progenitor type and promoting their neural stem cell properties.

### Conditional CA-YAP expression in embryonic mouse neocortex increases proliferative capacity of BPs

Next, we directly examined the potential effects of conditional CA-YAP expression in embryonic mouse neocortex on BP proliferation, using three distinct approaches. First, we performed immunofluorescence for the cycling cell marker Ki67 2 days (Figure 3A-C) and 3 days (Figure S4A,B) after IUE. Conditional CA-YAP expression doubled to tripled the proportion of Ki67-positive cells among the RFP-positive progeny of the targeted cells in the SVZ (Figure 3D, Figure S4C). Concomitant with this effect, conditional CA-YAP expression resulted in an altered distribution of the RFP-positive cells across the various zones of cortical wall, with a greater proportion of these cells in the VZ and a lesser proportion in the IZ and CP (Figure S5). These data raise the possibility of a relationship between the CA-YAP–induced progenitor proliferation and the reduced migration of the RFP-positive progeny of the targeted cells beyond the germinal zones to the basal region of the cortical wall.

**Figure 3.**
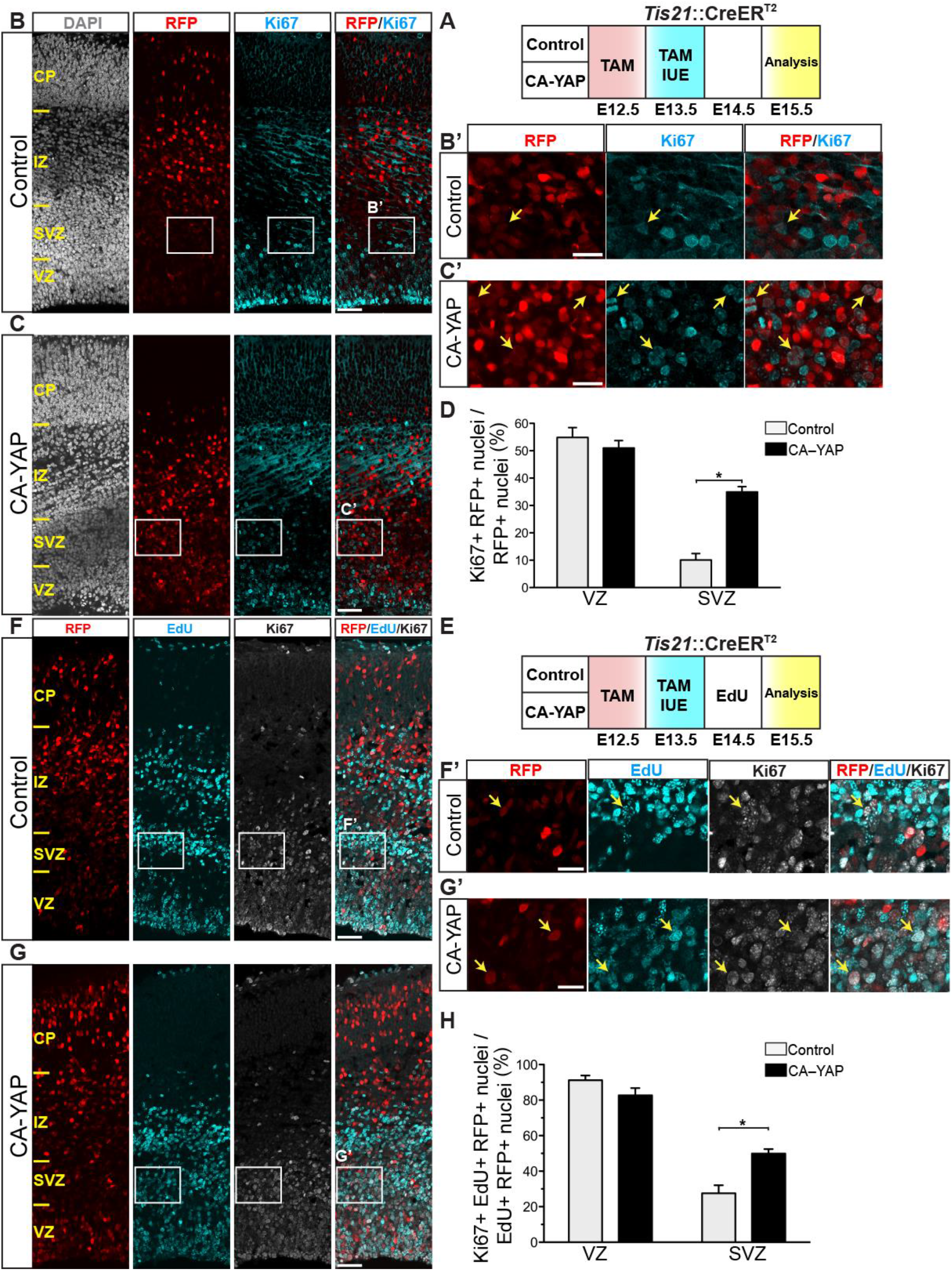
Conditional CA-YAP expression in the BP-genic lineage of embryonic mouse neocortex promotes BP proliferation and cell cycle re-entry. See also Figures S3, S4 and S5. *Tis21*::CreER^T2^ heterozygous mouse embryos received tamoxifen (TAM) at E12.5 and E13.5, the neocortex was *in utero* electroporated (IUE) at E13.5 with control plasmid (B, D, F, H) or CA-YAP–expressing plasmid (C, D, G, H) (see Fig. 2A), embryos did not (B-D) or did (F-H) receive a single EdU pulse at E14.5, and the neocortex was analyzed at E15.5, as shown in the flow schemes of the experiments in (A) and (E), respectively. (B, C) Double immunofluorescence for RFP (red) and Ki67 (cyan), combined with DAPI staining (white). Boxes indicate areas in the SVZ that are shown at higher magnification in panels B’ and C’; arrows indicate selected RFP-positive nuclei that are Ki67-positive. (D) Quantification of the percentage of RFP-positive nuclei that are Ki67-positive in the VZ and SVZ, upon control (light grey) and CA-YAP (black) electroporation. (F, G) Triple (immuno)fluorescence for RFP (red), EdU (cyan) and Ki67 (white). Boxes indicate areas in the SVZ that are shown at higher magnification in panels F’ and G’; arrows indicate selected RFP- and EdU-positive nuclei that are Ki67-positive. (H) Quantification of the percentage of RFP- and EdU-positive nuclei that are Ki67-positive in the VZ and SVZ, upon control (light grey) and CA-YAP (black) electroporation. (B, C, F, G) Images are 1-µm optical sections. Scale bars, 50 µm in (B, C, F, G), 20 µm in (B’, C’, F’, G’). (D, H) Two images (1-µm optical sections), each of a 200 µm-wide field of cortical wall, per embryo were taken, and the percentage values obtained were averaged for each embryo. Data are the mean of 4 embryos from four separate litters. The mean ± SEM is shown; * *P* <0.05 (Mann-Whitney *U*-test).

Second, we performed immunofluorescence for the mitotic cell marker phosphovimentin (pVIM). Conditional CA-YAP expression increased the proportion of pVIM-positive cells among the RFP-positive progeny of the targeted cells in the SVZ and (albeit not statistically significant) VZ (Figure S4D-F). Consistent with this, conditional CA-YAP expression increased the abundance of mitotic, pVIM-positive BPs and (albeit not statistically significant) APs (sum of RFP-positive and -negative mitoses) (Figure S4D,E,G).

Third, we carried out a cell-cycle re-entry assay. To this end, we performed a 5’-ethynyl-2’deoxyuridine (EdU) pulse labeling 24 h after IUE, followed by Ki67 immunofluorescence 24 h later (Figure 3E). Given the cell-cycle parameters of *Tis21*-GFP–positive APs and BPs at this stage of embryonic mouse neocortex development (Arai et al., 2011), the latter time interval is sufficient for the incorporated EdU to become inherited by daughter cells. Thus, EdU- and RFP-positive cells that are also positive for Ki67 are daughter cells that re-entered the cell-cycle (Figure 3E). Using this assay, we found that conditional CA-YAP expression doubled the cell-cycle re-entry of BPs in the SVZ (Figure 3, E-H). Taken together, the results of these three lines of investigation demonstrate that conditional CA-YAP expression in embryonic mouse neocortex increases BP proliferation.

### Conditional CA-YAP expression in embryonic mouse neocortex results in reduced deep-layer neuron and increased upper-layer neuron generation

Given that BPs generate most cortical neurons (Florio and Huttner, 2014; Lui et al., 2011), we explored the potential consequences of the CA-YAP–induced increase in BP proliferation for neuron generation in the embryonic mouse neocortex. To this end, we performed immunostaining for Tbr1, a deep-layer neuron marker, and Satb2, an upper-layer neuron marker 4 days after IUE (Figure 4A-E). Conditional CA-YAP expression reduced the proportion of Tbr1-positive neurons among the RFP-positive progeny of the targeted cells in the IZ and CP to almost half of control (29% vs. 44%, Figure 4F), and caused a small but statistically significant increase in the proportion of Satb2-positive neurons (from 76% to 87%, Figure 4G).

**Figure 4.**
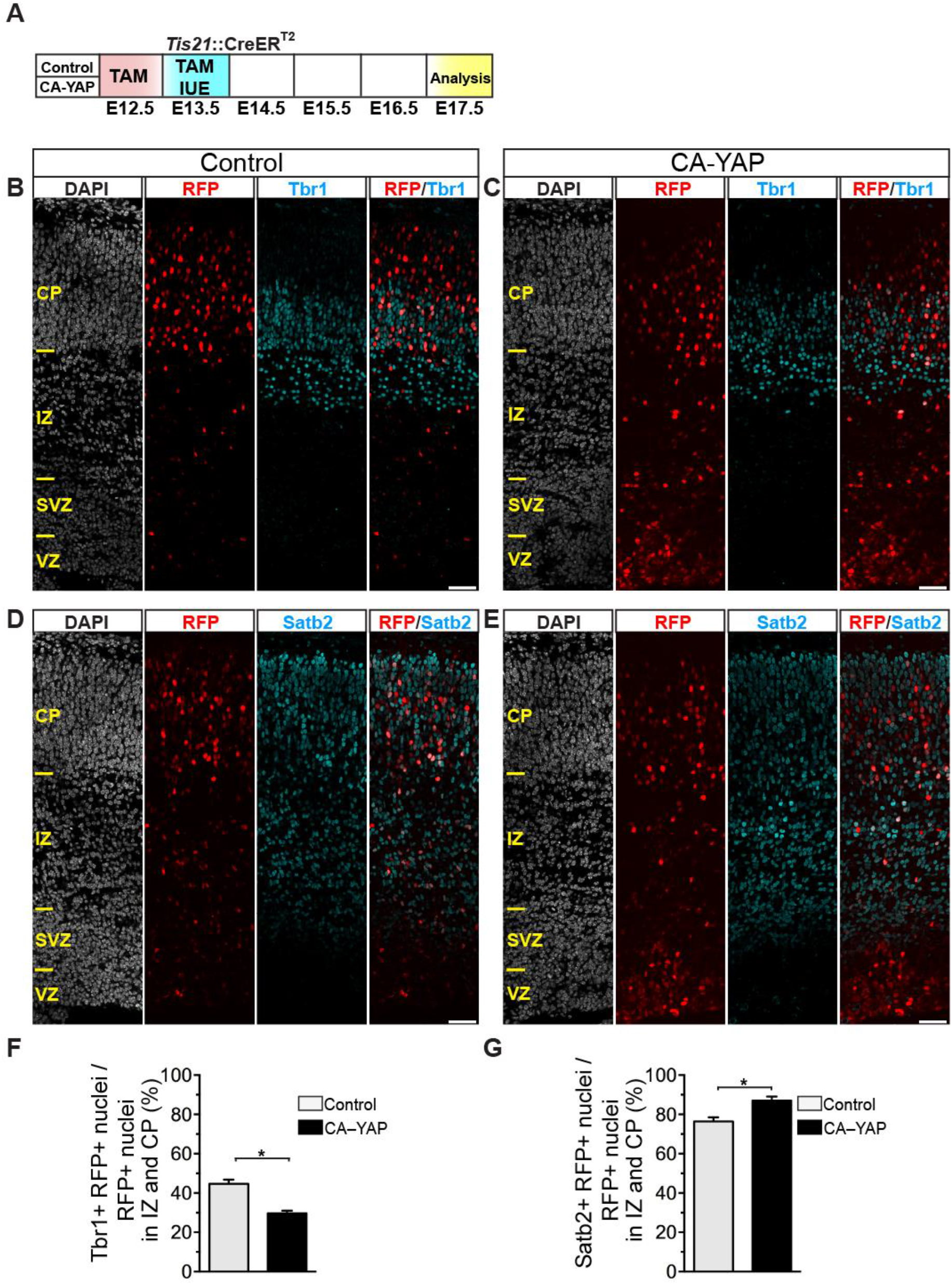
Conditional CA-YAP expression in the BP-genic lineage of embryonic mouse neocortex decreases production of deep-layer neurons and increases production of upper-layer neurons. See also Figure S6. (A) Flow scheme of experiments. *Tis21*::CreER^T2^ heterozygous mouse embryos received tamoxifen (TAM) at E12.5 and E13.5, and the neocortex was *in utero* electroporated (IUE) at E13.5 with control plasmid (B, D, F, G) or CA-YAP–expressing plasmid (C, E, F, G) (see Fig. 2A), followed by analysis at E17.5. (B-E) Double immunofluorescence for RFP (red) and either Tbr1 (B, C) or Satb2 (D, E) (cyan), combined with DAPI staining (white). Images are 1-µm optical sections. Scale bars, 50 µm. (F, G) Quantification of the percentage of RFP-positive nuclei that are Tbr1-positive (F) and Satb2-positive (G) in the intermediate zone (IZ) and cortical plate (CP), upon control (light grey) and CA-YAP (black) electroporation. Two images (1-µm optical sections), each of a 200 µm-wide field of cortical wall, per embryo were taken, and the percentage values obtained were averaged for each embryo. Data are the mean of 4 embryos from four separate litters. The mean ± SEM is shown; * *P* <0.05 (Mann-Whitney *U*-test).

Considering that the pool size of the RFP-positive progeny in the IZ and CP, four days upon IUE (E13-E17), is not much affected by the conditional CA-YAP expression compared to control (Figure S6), these data are consistent with the notion that the changes in the proportions of specific types of neurons among the RFP-positive progeny reflect the changes in neuron generation in embryonic mouse neocortex. Specifically, on the one hand, the CAYAP–induced increase in BP proliferation initially results in a reduced production of deep-layer neurons because upon conditional CA-YAP expression a certain proportion of the mouse BPs has been induced to undergo symmetric proliferative divisions and hence a lesser proportion of the mouse BPs is available to undergo the symmetric consumptive divisions that generate neurons. On the other hand, the CA-YAP–induced increase in BP proliferation eventually results in an increased mouse BP pool size that at later stages of cortical neurogenesis gives rise to more upper-layer neurons.

### Pharmacological inhibition of YAP activity reduces mitotic BP abundance in embryonic ferret and fetal human neocortex

Having established that mimicking a ferret- or human-like expression of active YAP in BPs of embryonic mouse (i.e. lissencephalic) neocortex suffices to increase their proliferation, we next investigated whether the presence of active YAP in BPs of developing ferret and human (i.e. gyrencephalic) neocortex is a necessary requirement for their proliferation. To this end, we applied an inhibitor of YAP, verteporfin (Brodowska et al., 2014; Liu-Chittenden et al., 2012; Song et al., 2014; Wang et al., 2016), in an *ex vivo* free-floating tissue (FFT) culture system (Schenk et al., 2009), using E33-34 ferret and 11-13 wpc human neocortical tissue (Figure 5). Verteporfin is a small molecule that inhibits the association of YAP with TEAD transcription factors and thereby prevents the expression of genes linked to cell proliferation (Liu-Chittenden et al., 2012).

**Figure 5.**
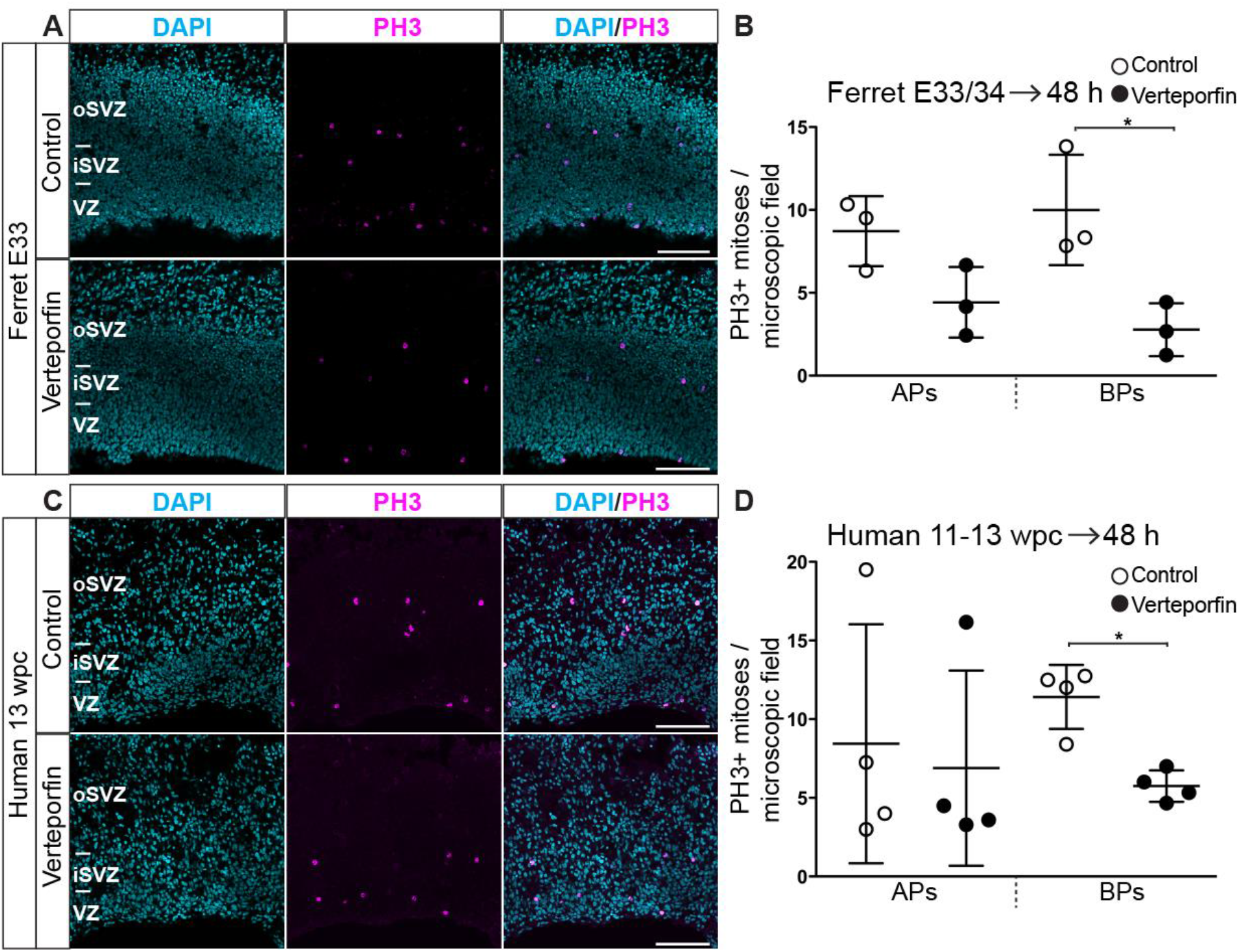
Inhibition of YAP activity by verteporfin reduces mitotic BP levels in embryonic ferret and fetal human neocortex. Ferret E33-34 (A, B) and human 11-13 wpc (C, D) neocortex was incubated for 48 h in FFT culture, in the absence (Control, upper rows in (A, C), open circles in (B, D)) or presence (lower rows in (A, C), filled circles in (B, D)) of 1 µM verteporfin, followed by analysis. (A, C) Immunofluorescence for PH3 (magenta), combined with DAPI staining (blue), upon control (upper rows) and verteporfin (lower rows) treatment. Images are 1-µm optical sections. Scale bars, 100 µm. (B, D) Quantification of the number of APs and BPs in mitosis, as revealed by PH3 immunofluorescence, per microscopic field (400 µm-wide field of cortical wall), upon control (open circles) and verteporfin (filled circles) treatment. Five to eight images (1-µm optical sections), per either ferret embryo (B) or human fetus (D) were taken, and the values obtained were averaged for each embryo/fetus. Data are the mean of 3 ferret embryos from three separate litters (B) and of 4 human fetuses (D). Error bars indicate SD; * *P* <0.05 (Mann-Whitney *U*-test).

Treatment of embryonic ferret and fetal human neocortical FFT cultures with 1 µM verteporfin for 48 h decreased the abundance of basal mitoses, identified by phosphohistone H3 (PH3) immunofluorescence (Figure 5A,C), approximately 4-fold and 2-fold, respectively (Figure 5B,D). In ferret, verteporfin treatment also resulted in a decrease, albeit not statistically significant, in apical mitoses (Figure 5B).

### Dominant-negative YAP expression reduces mitotic BP abundance in embryonic ferret neocortex

We sought to obtain corroborating in vivo evidence for an essential role of YAP activity in BP proliferation in developing gyrencephalic neocortex. To this end, we used a dominant-negative YAP construct (DN-YAP) to block its transcriptional co-activator function (Nishioka et al., 2009; Sudol et al., 2012) (see Supplemental Experimental Procedures for details). In this DN-YAP construct, the transactivation domain of mouse YAP is replaced with the engrailed domain, a *Drosophila* repressor of transcription (Nishioka et al., 2009), and nuclear localization of DN-YAP is ensured by replacing YAP serine112 with alanine (Hao et al., 2007; Zhao et al., 2007). Forced expression of DN-YAP creates multiple copies of DN-YAP that outcompete endogenous wtYAP in binding to TEAD transcription factors, which causes downregulation of YAP-driven genes linked to proliferation (Nishioka et al., 2009; Zanconato et al., 2015). We chose embryonic ferret neocortex to examine the effects of DN-YAP expression on BP proliferation.

We performed IUE of ferret embryos to deliver the DN-YAP construct into the developing ferret neocortex (Kawasaki et al., 2012; Kawasaki et al., 2013). Specifically, we coelectroporated the ferret dorsolateral neocortex with either CAGGS-empty vector plus CAGGS-EGFP or with CAGGS-DN-YAP plus CAGGS-EGFP. IUE was performed at E33, the stage which corresponds to mid-neurogenesis in the ferret. Ferret embryos were harvested 2 days later, at E35, to allow for sufficient time for the targeted APs to complete their cell-cycle (Turrero Garcia et al., 2015) and give rise of BPs (Figure 6A). To confirm the expression of DN-YAP, we performed immunostaining using a YAP antibody that recognizes both endogenous YAP and DN-YAP and compared the level of YAP immunoreactivity in GFP-positive cells upon DN-YAP expression to that of control. Upon DN-YAP expression, many of the GFP-positive cells showed a higher level of YAP immunoreactivity, especially in the SVZ, suggesting that the DN-YAP was successfully expressed in BPs (Figure 6B, compare lower and upper rows).

**Figure 6.**
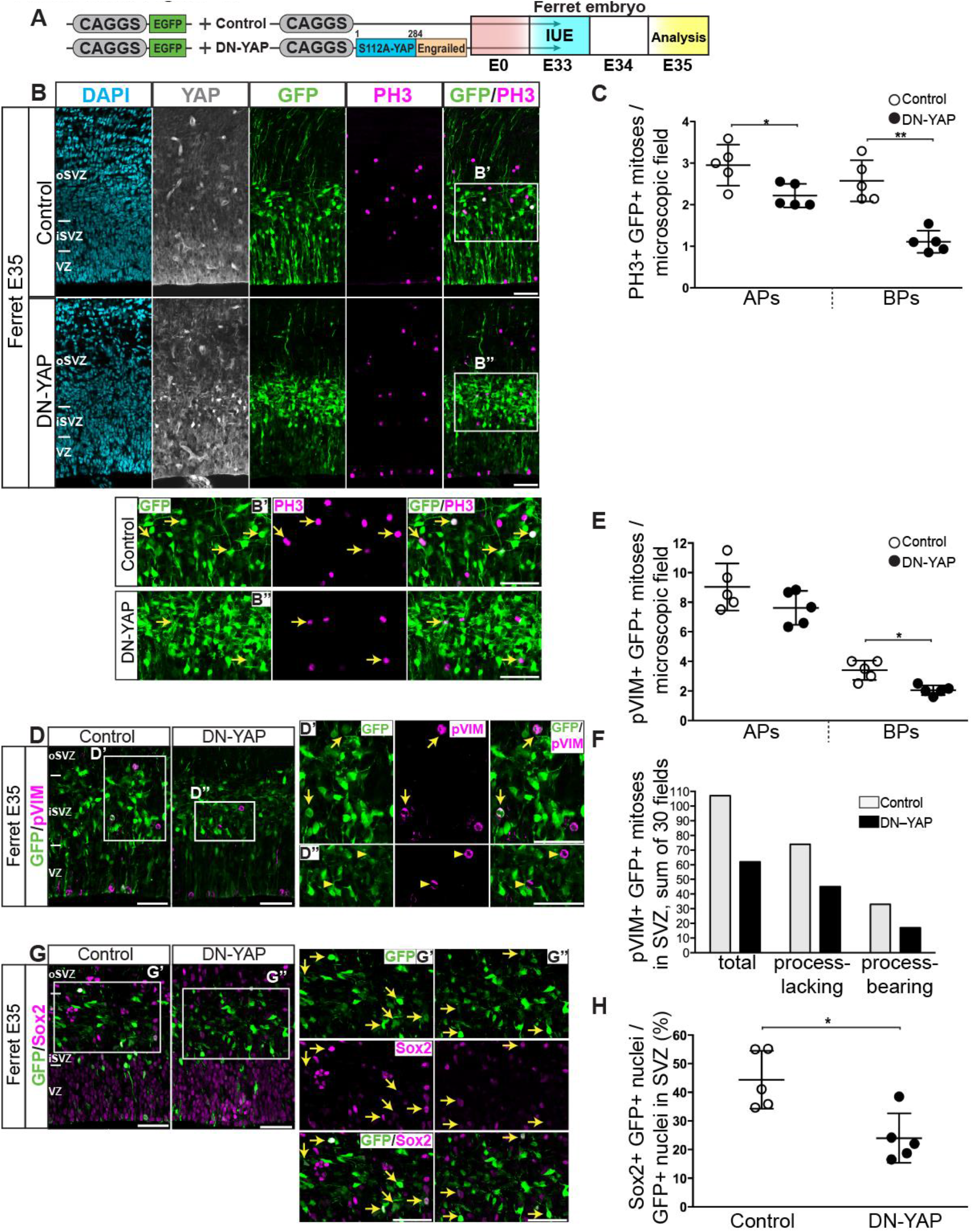
Expression of a DN-YAP in embryonic ferret neocortex reduces mitotic BP levels. (A) Left, cartoon showing the EGFP-expressing plasmid plus the control plasmid (top) and the EGFP-expressing plasmid plus the DN-YAP–expressing plasmid (bottom). Right, flow scheme of experiment. Ferret neocortex was electroporated at E33 with either GFP-expressing plus control plasmids (B upper row, B’, C, D left, D’, E, F, G left, G’, H) or GFP-expressing plus DN-YAP–expressing plasmids (B lower row, B’’, C, D right, D’’, E, F, G right, G’’, H), followed by analysis at E35. (B) Triple immunofluorescence for YAP (white), GFP (green) and PH3 (magenta), combined with DAPI staining (blue). Boxes indicate areas in the SVZ that are shown at higher magnification in panels B’ and B”; arrows indicate selected GFP-positive cells that are PH3-positive. (C) Quantification of the number of GFP-positive APs and BPs in mitosis, as revealed by PH3 immunofluorescence, per microscopic field (200 µm-wide field of cortical wall), upon control (open circles) and DN-YAP (filled circles) electroporation. Six to fifteen images (1-µm optical sections) per ferret embryo were taken, and the values obtained were averaged for each embryo. The data show the averaged values for 5 ferret embryos from one litter. (D) Double immunofluorescence for GFP (green) and pVIM (magenta). Boxes indicate areas in the SVZ that are shown at higher magnification in panels D’ and D”; arrows in (D’) indicate selected GFP-positive cells that are pVIM-positive; arrowheads in (D”) indicate selected pVIM-positive nuclei that are GFP-negative. (E) Quantification of the number GFP-positive APs and BPs in mitosis, as revealed by pVIM immunofluorescence, per microscopic field (200 µm-wide field of cortical wall), upon control (open circles) and DN-YAP (filled circles) electroporation. Six images (each a 5-µm z-stack) per ferret embryo were taken, and the values obtained were averaged for each embryo. The data show the averaged values for 5 ferret embryos from one litter. (F) Quantification of the number of total (left), of process-lacking (middle) and of process-bearing (right) GFP-positive mitoses in the SVZ, as revealed by pVIM immunofluorescence, upon control (light grey) and DN-YAP (black) electroporation. Six images (each a 5-µm z-stack of a 200 µm-wide field of cortical wall) per ferret embryo were taken, 5 ferret embryos from one litter were analyzed, and data are the sum of the thirty fields counted. (G) Double immunofluorescence for GFP (green) and Sox2 (G) (magenta). Boxes indicate areas in the SVZ that are shown at higher magnification in panels G’ and G”; arrows indicate selected GFP-positive nuclei that are Sox2-positive. (H) Quantification of the percentage of GFP-positive cells in the SVZ that are Sox2-positive, upon control (open circles) and DN-YAP (filled circles) electroporation. Six images (1-µm optical sections), each of a 200 µm-wide field of cortical wall, were taken per ferret embryo, and the percentage values obtained were averaged for each embryo. Data are the mean of 5 ferret embryos from one litter. (B, B’, B”, D, D’, D”, G, G’, G”) Images are 1-µm optical sections. Scale bars, 50 µm. (C, E, H) The mean ± SD is shown; * *P* <0.05, ** *P* <0.01 (Mann-Whitney *U*-test).

Analysis of the GFP-positive progeny of the targeted APs revealed that expression of DN-YAP decreased slightly the abundance of mitotic APs, and massively that of mitotic BPs, as revealed by PH3 immunofluorescence (Figure 6B,C). Similar results were obtained by quantitation of pVIM-positive mitoses (Figure 6D,E).

There are two main types of BPs, (i) those lacking processes at mitosis, i.e. bIPs, and (ii) those bearing radial processes at mitosis, i.e. bRG (Fietz et al., 2010; Kelava et al., 2012; Reillo et al., 2011). To examine whether any of these two types of BPs is preferentially affected by DN-YAP expression, we analyzed the processes as revealed by pVIM staining in GFP-positive mitotic BPs in the SVZ. We observed a similar decrease in process-lacking and process-bearing BPs, which suggests that the effect of DN-YAP expression is non-selective with regard to the BP population type, affecting equally bIP and bRG (Figure 6F).

Given that conditional CA-YAP expression in embryonic mouse neocortex resulted in a specific increase in Sox2-positive BPs (Figure 2G), we explored whether inhibition of YAP activity in embryonic ferret neocortex would affect the Sox2-positive pool of BPs in the SVZ. Indeed, DN-YAP expression reduced the proportion of Sox2-positive BPs among the GFP-positive progeny of the targeted cells in the SVZ (Figure 6G,H).

Taken together, our data using verteporfin and DN-YAP indicate that YAP activity is necessary for the proliferation of BPs in developing gyrencephalic neocortex *in vivo*.

## Discussion

The present study demonstrates a crucial role of YAP, the central effector of the Hippo signaling pathway, in the development and evolution of the mammalian neocortex. Specifically, our findings advance our understanding of cNPC activity in cortical development, and of its differences across mammals, with regard to two key aspects. First, YAP activity is shown to be necessary and sufficient to maintain the proliferative capacity of BPs at levels typically seen in developing gyrencephalic neocortex. Second, differences in YAP activity in BPs between embryonic mouse, which develops a lissencephalic neocortex, on the one hand, and embryonic ferret and fetal human, which develop a gyrencephalic neocortex, on the other hand, are shown to contribute to, if not underlie, the differences in BP proliferative capacity across these species. Three aspects of our findings deserve particular consideration.

### DIFFERENTIAL YAP EXPRESSION BETWEEN VZ AND SVZ ACROSS SPECIES

Previous reports on the role of YAP were confined to mouse brain development and concentrated on APs (Cappello et al., 2013; Lavado et al., 2013; Lavado et al., 2014; Saito et al., 2017). In contrast to these reports, the present study is focused on BPs and in this context has compared YAP expression and activity in the SVZ vs. VZ of developing neocortex of three mammals exhibiting a different extent of neocortex expansion – mouse, ferret and human. Our demonstration that in species in which BPs exhibit significant proliferative capacity, YAP expression and activity are well detectable not only in the VZ but also SVZ, has at least two implications. First, to the best of our knowledge, these data constitute the first report of YAP protein expression in the iSVZ and oSVZ of developing neocortex. Second, the present findings suggest that the role of YAP, and hence of Hippo signaling, in the proliferation and consequently the pool size of cNPCs is far more widespread than previously assumed.

### YAP IN BPS – A PUTATIVE DOWNSTREAM TARGET OF SOX2

The co-expression of active YAP with Sox2 in ferret and human BPs suggests an interesting mechanistic scenario regarding the regulation of YAP expression and activity. Recently, it was shown that Sox2 directly drives the expression of YAP in progenitors of the osteo-adipo lineage (Seo et al., 2013). Our results therefore raise the possibility that YAP expression in proliferative BPs is driven by Sox2. Furthermore, while YAP activity is known to be decreased by phosphorylation via upstream Hippo signaling kinases such as Nf2 and Wwc1 (Yu et al., 2015), it was recently reported that in cancer stem cells Sox2 represses transcription of the *NF2* and *WWC1* genes, which in turn increased YAP activity (Basu-Roy et al., 2015). Our finding of increased YAP activity in proliferative BPs of developing gyrencephalic neocortex could therefore be explained by Sox2 in these cells repressing *NF2* and *WWC1*.

### BP FATE SWITCH UPON CONDITIONAL YAP EXPRESSION

However, our data are not only consistent with the notion that in ferret and human BPs Sox2 positively regulates YAP expression levels and activity, but also indicate that forced YAP expression in mouse BPs, directly or indirectly, results in increased Sox2 expression and decreased Tbr2 expression. In other words, the present approach of conditionally expressing CA-YAP in the mouse Tis21-positive BP lineage induced a BP fate switch from neurogenic to proliferative. This in turn led to an expansion of the BP pool, which eventually resulted in an increased generation of upper-layer neurons. Hence, increasing YAP activity in BPs is sufficient to induce features that are hallmarks of an expanded neocortex, as characteristically observed in gyrencephalic mammals (Florio and Huttner, 2014; Lui et al., 2011). This, together with our finding that inhibiting YAP activity in BPs of developing ferret and human neocortex reduced their abundance, in turn leads us to conclude that an increased YAP activity in BPs of developing neocortex is a major contributor to its evolutionary expansion.

## Experimental Procedures

### Ethics

All animal experiments (mice and ferrets) were performed in accordance with the German Animal Welfare legislation (“Tierschutzgesetz”). All procedures regarding the animal experiments were approved by the Governmental IACUC (“Landesdirektion Sachsen”) and overseen by the Institutional Animal Welfare Officer(s). The license numbers concerning the experiments with mice are: Untersuchungen zur Neurogenese in Mäuseembryonen TVV2015/05 (*in utero* electroporation, tamoxifen, EdU) and 24–9168.24-9/2012-1 (tissue collection without prior *in vivo* experimentation). The license number concerning the experiments with ferrets is: Untersuchungen zur Neurogenese in Frettchen" (TVV 2015/02) issued by “Landesdirektion Sachsen”.

### Mice

To characterize YAP expression E13-14 mouse embryos (C57BL/6JOlaHsd) were used. For electroporation E13 *Tis21*-CreER^T2+/-^ heterozygous mouse embryos were used. These were obtained by crossing C57BL/6JOlaHsd females and *Tis21*-CreER^T2+/+^ males (Wong et al.,2015). Mice were crossed and kept under strict pathogen-free conditions in the animal facility of the Max Planck Institute of Molecular Cell Biology and Genetics.

### Ferrets

Normally pigmented pregnant female sable ferrets (*Mustela putorius furo*) were purchased from Marshall BioResources (North Rose, NY, USA). They were delivered and housed in the animal facility of the Max Planck Institute of Molecular Cell Biology and Genetics.

### Human fetal tissue

Human fetal brain tissue was obtained from two sources. First, from the Klinik und Poliklinik für Frauenheilkunde und Geburtshilfe, Universitätsklinikum Carl Gustav Carus of the Technische Universität Dresden, following elective pregnancy termination and informed written maternal consents, and with approval of the local University Hospital Ethical Review Committees. The age of a 12 wpc fetus (n=1) was assessed by ultrasound measurements of crown-rump length and other standard criteria of developmental stage determination. The second source was the Human Developmental Biology Resource (HDBR). This human fetal brain tissue was provided by the Joint MRC/Wellcome Trust (grant # MR/R006237/1) Human Developmental Biology Resource (www.hdbr.org). The HDBR provided fresh tissue from fetuses aged 11-13 wpc, (11 wpc, n = 4; 12 wpc, n = 2; 13 wpc, n = 2; wpc 14, n=1). Human fetal brain tissue was dissected in 1x PBS and used immediately for culture or fixation (as indicated) when obtained from Dresden. When obtained from HDBR, tissue was dissected and shipped in Hibernate E media (Gibco A1247601). Upon arrival, all tissue was cultured in slice culture medium (SCM, see section on *ex vivo* FFT cultures) for 2-3 h prior to any further manipulation. All tissue was fixed for at least 24 h at 4°C in 4% paraformaldehyde in 120 mM phosphate buffer (pH 7.4) (referred to in short as 4% PFA).

### Tamoxifen preparation and administration

Tamoxifen powder, 200 mg, was dissolved in 10 ml of corn oil (Sigma, T-5648) under constant stirring at ≈40°C. Tamoxifen was administered to trigger activation of Cre recombinase in *Tis21*::CreER^T2 +/-^ mouse embryos. Pregnant mice received tamoxifen (2 mg, 0.1 ml) orally by gavage at E12.5, i.e. one day before IUE, and once on the day of IUE (E13.5).

### *In utero* electroporation of mice

Tamoxifen-treated pregnant mice carrying E13.5 embryos were anesthetized using initially 5% isoflurane (Baxter, HDG9623), followed by 2-3% isoflurane during the IUE procedure. Endotoxin-free plasmids (Control, *pCAGGS–LoxP–Gap43-GFP–LoxP–IRES–nRFP*; CA-YAP, *pCAGGS–LoxP–Gap43-GFP–LoxP–CA-YAP–IRES–nRFP*) were mixed on the day of surgery with Fast Green (Sigma, 0.25% final concentration) to a final plasmid concentration of 2 µg/µl in 1x PBS. Using a borosilicate microcapillary (Sutter instruments, BF120-69-10) the DNA/Fast Green mixture was intraventricularly injected, which was followed by six 50-msec pulses of 30 V at 1 sec intervals (BTX genetronics Inc., 45-0052INT), using a 3-mm diameter electrode (BTX genetronics Inc., 45-0487). After the IUE, the uterus was placed back into the abdominal cavity, and the peritoneum was sutured (VICRYL™ 5-0, V493H). Abdominal skin was closed with clips and animal received 100 µl of painkiller (Rimadyl 1 mg/ml). Pregnant mice were sacrificed by cervical dislocation at the indicated time points (E14.5-E17.5), and embryonic brains were dissected and fixed in 4% PFA, overnight at 4°C.

### Verteporfin treatment of ferret and human neocortex in *ex vivo* free-floating tissue culture

An *ex vivo* free-floating tissue (FFT) culture system, adapted and modified from (Schenk et al., 2009; Long et al., 2018), was used to perform verteporfin treatment of embryonic ferret and fetal human neocortex tissue. E33 ferret brains were dissected, meninges removed, and the two hemispheres separated. Fetal human neocortex tissue of 11-13 wpc was cut into 2000-2500 µm-thick pieces (tangential dimension). Tissue was cultured in a whole-embryo culture incubator (Ikemoto RKI) in a rotating flask with 1.5 ml of SCM (for composition, see below), and incubated at 37°C in the presence of a humidified atmosphere consisting of 40% O_2_ / 5% CO_2_ / 55% N_2_, with continuous rotation at 26 rpm. Control flasks contained 10 µl of DMSO (Dimethyl sulfoxide, Sigma, 472301) added to the 1.5 ml of SCM. Verteporfin flasks contained 1 µM of verteporfin (Verteporfin, Sigma, SML0534) dissolved in 10 µl of DMSO, added to the 1.5 ml of SCM. FFT cultures were carried out for 48 h, with one change of SCM (containing either DMSO or DMSO plus verteporfin) after 24 h. After 48 h of FFT culture, tissue was fixed in 4% PFA overnight at 4°C.

The SCM medium used for FFT cultures contained: 84 ml of Neurobasal medium (Gibco, 21103049) supplemented with either 10 ml of rat serum (ferret and human cultures) or 10 ml of 5x KnockOUT^TM^ Serum Replacement (for human cultures, Gibco, 10828028), 1 ml GlutaMAX^TM^ (100x), 1 ml penicilin/streptomycin (100x) (Gibco,15140122), 1 ml N-2 (100x) (Gibco, 17502048), 2 ml B-27 (50x) (Gibco, 17504044) and 1 ml of 1 M HEPES-NaOH, pH 7.2, to yield a final volume of 100 ml.

### *In utero* electroporation of ferrets

*In utero* electroporation of E33 ferret embryos was performed as originally established (Kawasaki et al., 2012; Kawasaki et al., 2013), with the modifications indicated below. Pregnant ferrets (with embryos at E33) were kept fasted for at least 3 h before the surgery and placed in the narcosis box with 4% isoflurane. Subsequently, they were positioned on the operation table and attached to the narcosis mask with 3% isoflurane and injected subcutaneously with analgesic (0.1 ml Metamizol, 50 mg/kg), antibiotic (0.13 ml Synulox, 20 mg/kg or 0.1 ml amoxicilin, 10 mg/kg) and glucose (10 ml 5% glucose solution). The ferret bellies were then shaved, sterilized with iodide and surgically opened. Then, the uterus was exposed. As the ferret uterus is pigmented, a transmitted light source was used for the visualization of embryos. Embryos were injected intraventricularly with a solution containing 0.1% Fast Green (Sigma) in sterile 1x PBS, 2 µg/µl of one of the endotoxin-free plasmids (CTRL-empty or DN-YAP) as indicated. To visualize electroporated cells, CTRL and DN-YAP plasmids were co-electroporated with CAGGS-EGFP (1 µg/µl), using a 5-mm diameter electrode (BTX genetronics Inc., 45-0489). Electroporations were performed with six 50-msec pulses of 100 V at 1 sec intervals. Subsequently, the uterus was placed back in the peritoneal cavity, muscle layer with the peritoneum were sutured (VICRYL™ 4-0), after which the skin was sutured intracutaneously. Animals were carefully monitored until they woke up and then underwent postoperative care for the following 3 days. Pregnant females received subcutaneous injections of 15 ml of 5% glucose and the painkiller Metamizol (50 mg/kg, WDT, 99012) three times per day.

The embryos were obtained by cesarean section at E35, and the cerebral cortex was dissected and fixed in 4% PFA overnight at 4°C. To this end, the mother ferrets underwent a second surgery that followed the same pre-operative care, anesthesia and analgesia as the first surgery. The sutures from the first operation were removed and the uterus exposed, after which the embryos were removed by a caesarian section. Subsequently a complete hysterectomy was performed, after which the muscle layer with peritoneum and skin were sutured and the animal underwent the same post-operative care as after the first surgery. Animals were kept at the BMS of the MPI-CBG for at least two weeks after the second surgery after which they were donated for adoption.

### Statistical analysis

Data were tabulated in Excel (Microsoft, Redmond, WA) and analyzed in Prism 5 (GraphPad) software. Statistical analyses were performed using unpaired Student’s *t*-test, Mann-Whitney *U*-test, and one-way ANOVA test.

## Acknowledgements

We are grateful to the Services and Facilities of the Max Planck Institute of Molecular Cell Biology and Genetics for the outstanding support provided, notably J. Helppi and his team of the Biomedical services (BMS), and J. Peychl and his team of the Light Microscopy Facility. We also thank J. Fei (Tanaka Lab, CRTD, Dresden, Germany) for providing YAP antibody and constructive suggestions. We would like to thank all members of the Huttner group for helpful discussions, especially A. Güven for advice, A. Sykes for assistance with molecular cloning and M. Florio for sharing RNA-seq data on YAP expression. We thank D. Gerrelli, S. Lisgo and their teams at the HDBR for the invaluable support from this resource. M.K. was a member of the International Max Planck Research School for Cell, Developmental and Systems Biology, and a doctoral student at the Technische Universität Dresden. W.B.H. was supported by grants from the DFG (SFB 655, A2) and the ERC (250197), by the DFG-funded Center for Regenerative Therapies Dresden and by the Fonds der Chemischen Industrie.

## Author contributions

M.K., T.N., J.T.L.M.P., and W.B.H. conceived the project and designed the experiments.

M.K. performed most of the experiments, with technical assistance from J.T.L.M.P., T.N., K.L. and N.K.

B.L. performed the cesarean sections on ferrets and took care of ferrets pre- and post-surgery.

H.K. shared the technical knowledge on IUE of ferrets.

M.K. analyzed the data, with day-to-day supervision by J.T.L.M.P. and T.N.

M.K. and W.B.H. wrote the manuscript, with input from T.N., K.L. and N.K.

W.B.H. supervised the project.

## Supplemental Information

1. Supplemental figures S1 to S6 with legends
2. Supplemental results
3. Supplemental experimental procedures
4. Supplemental references

**Figure S1.**
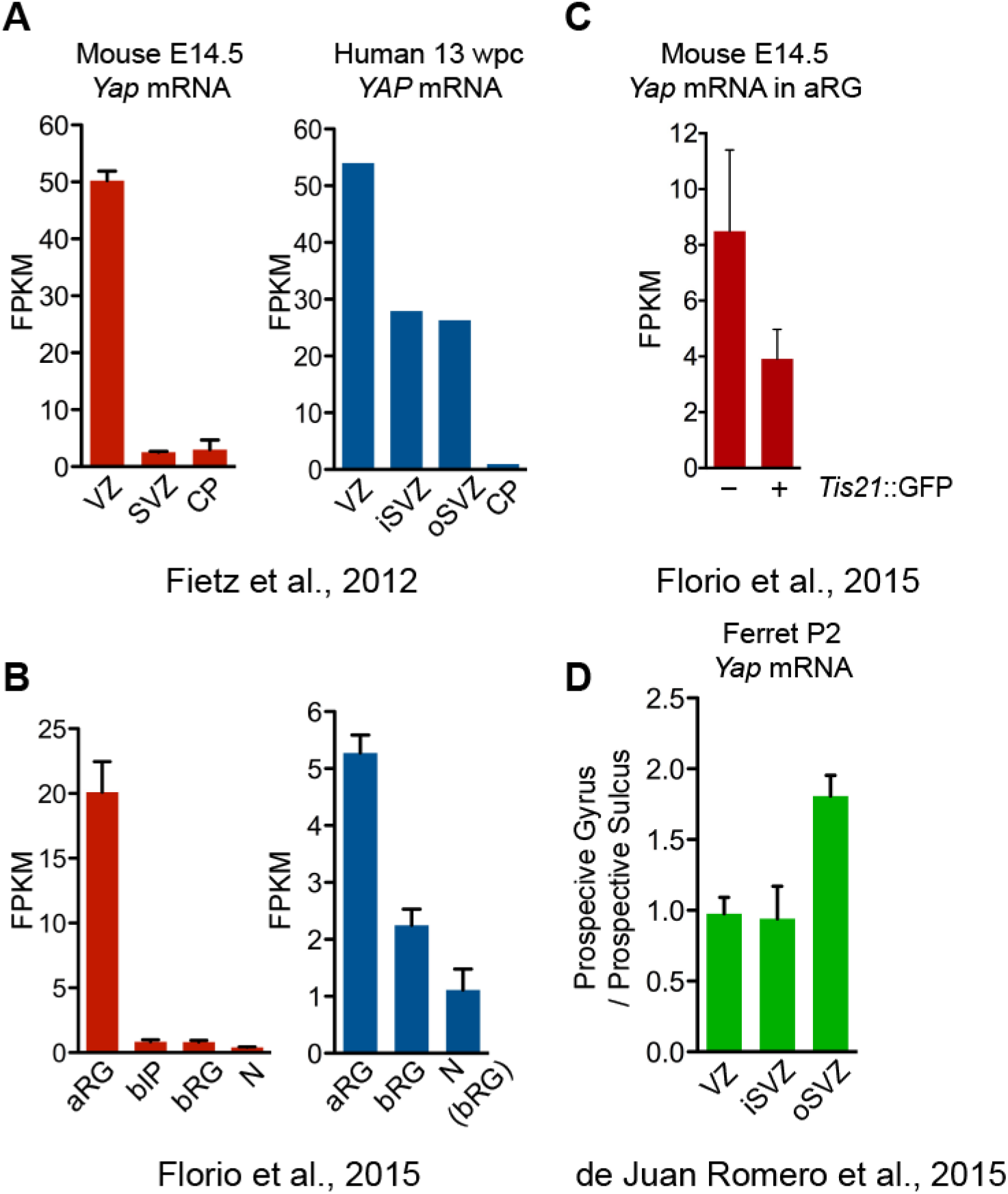
Fetal human and postnatal ferret, but not embryonic mouse, neocortical BPs express *YAP* mRNA. Related to Figure 1. (A, B) FPKM values of *Yap*/*YAP* mRNA in the mouse E14.5 (left) and human 13 wpc (right) neocortical germinal zones (A, determined in Fietz et al. 2012) and cNPC subpopulations (B, determined in Florio et al. 2015); bIP, mouse cell fraction containing bIPs and other prominin-1 and DiI double-negative cell bodies; N, mouse neurons; N(bRG), human neuron fraction containing bRG in G1 (see Florio et al. 2015). (C) FPKM values of *Yap* mRNA in mouse E14.5 *Tis21*::GFP-positive and -negative neocortical aRG (determined in Florio et al. 2015). (D) Ratio of *Yap* mRNA levels in prospective gyrus / sulcus in ferret postnatal day 2 (P2) neocortex (determined in de Juan Romero et al. 2015). Data are from one human transcriptome (A) or are the mean of 5 (A, C) and 4 (B) mouse transcriptomes, 4 (B) human transcriptomes, and 4 ferret transcriptomes (D); error bars indicate SD.

**Figure S2.**
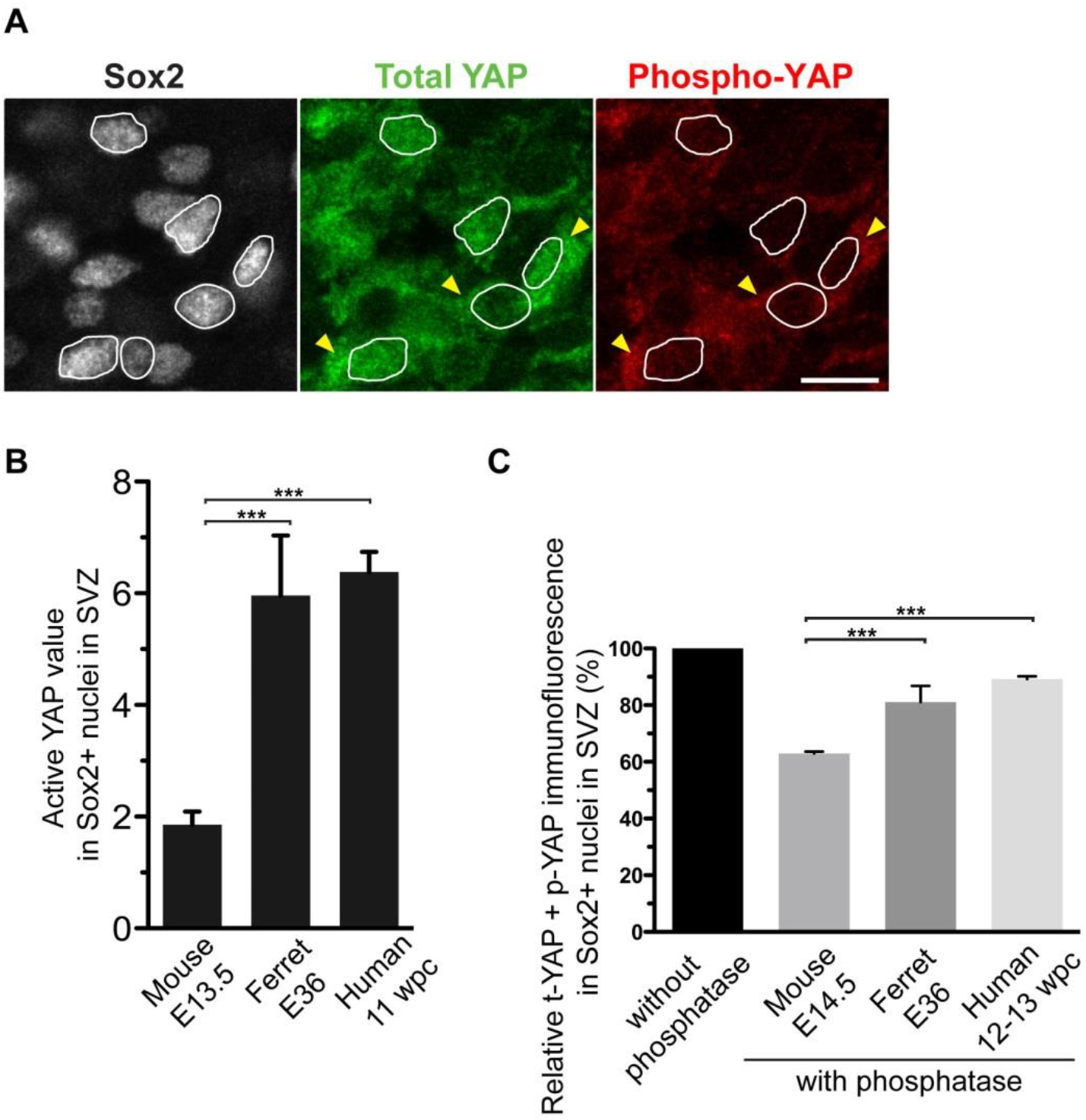
A greater proportion of the nuclear YAP in BPs is dephosphorylated at serine 127, i.e. active, in embryonic ferret and fetal human than embryonic mouse neocortex. Related to Figure 1. (A) Triple immunofluorescence for Sox2 (white), total YAP (i.e., YAP irrespective of phosphorylation; green) and phospho-YAP (P-serine 127; red) on a cryosection of human 11 wpc neocortex; the SVZ is shown. Selected Sox2 and YAP (green) double-positive BP nuclei are outlined by white lines; note the barely detectable levels of phospho-YAP compared to total YAP immunoreactivity in these nuclei compared to the cytoplasmic signals for total YAP and phospho-YAP (arrowheads). Scale bar, 10 µm. (B) Quantitation of active YAP levels in Sox2-positive BP nuclei in the SVZ of mouse E13.5, ferret E36 and human 11 wpc neocortex by comparison of total YAP and phospho-YAP immunofluorescence. Cryosections were subjected to triple immunofluorescence for Sox2, total YAP and phospho-YAP as described in panel A for human neocortex. One to two images per embryo/fetus (1 image per cryosection) were taken, and 30 randomly picked Sox2-positive nuclei in the SVZ were scored per image, as follows. First, for each image, the mean immunofluorescence intensity values for total YAP and for phospho-YAP from three representative areas of cytoplasm (see arrowheads in A) were determined, to serve as internal standards for the comparison of nuclear total YAP and nuclear phospho-YAP. The ratio of these two values was used to adjust the immunofluorescence intensity values for nuclear phospho-YAP relative to the immunofluorescence intensity values for nuclear total YAP. Then, for each nucleus, the adjusted immunofluorescence intensity value for phospho-YAP was subtracted from the immunofluorescence intensity value for total YAP, to yield the value for dephosphorylated, i.e. active, YAP. The values obtained for nuclear active YAP were averaged for each embryo/fetus. Data are the mean of 7 mouse, 4 ferret and 4 human embryos/fetuses. Error bars indicate SD; *** *P* <0.001 (one-way ANOVA test, post-hoc Tukey HSD). (C) Quantitation of the proportion of inactive YAP levels in Sox2-positive BP nuclei in the SVZ of mouse E14.5, ferret E36 and human 12-13 wpc neocortex by determining the effect of protein phosphatase treatment on the total YAP plus phospho-YAP immunofluorescence signal. Cryosections were treated without (control) or with (phosphatase treatment) lambda protein phosphatase, followed by double immunofluorescence for Sox2 and YAP. For YAP immunofluorescence, two rabbit monoclonal antibodies were used together, one recognizing YAP irrespective of serine127 phosphorylation (total YAP, t-YAP) and the other recognizing the serine127 phosphorylation site when phosphorylated (phospho-YAP, p-YAP). For each neocortex sample per species, three cryosections without and three cryosections with lambda protein phosphatase pre-treatment were analyzed; for each cryosection, the immunofluorescence intensity values obtained with the sum of these two antibodies were measured in 30 randomly selected Sox2+ BP nuclei in the SVZ, and the average value per cryosection was determined. For each species, the mean of these average values for the three control cryosections is set to 100% (black column), and the mean of these average values for the three phosphatase-treated cryosections is expressed relative to this. Data are the mean of three neocortex samples per species; error bars indicate SD; ** *P* <0.01, *** *P* <0.001 (one-way ANOVA test, post-hoc Tukey HSD). Note that the reduction, upon protein phosphatase treatment, in the YAP immunofluorescence signal obtained with the sum of the two antibodies (total YAP plus phospho-YAP) indicates the contribution of phospho-YAP to this signal.

**Figure S3.**
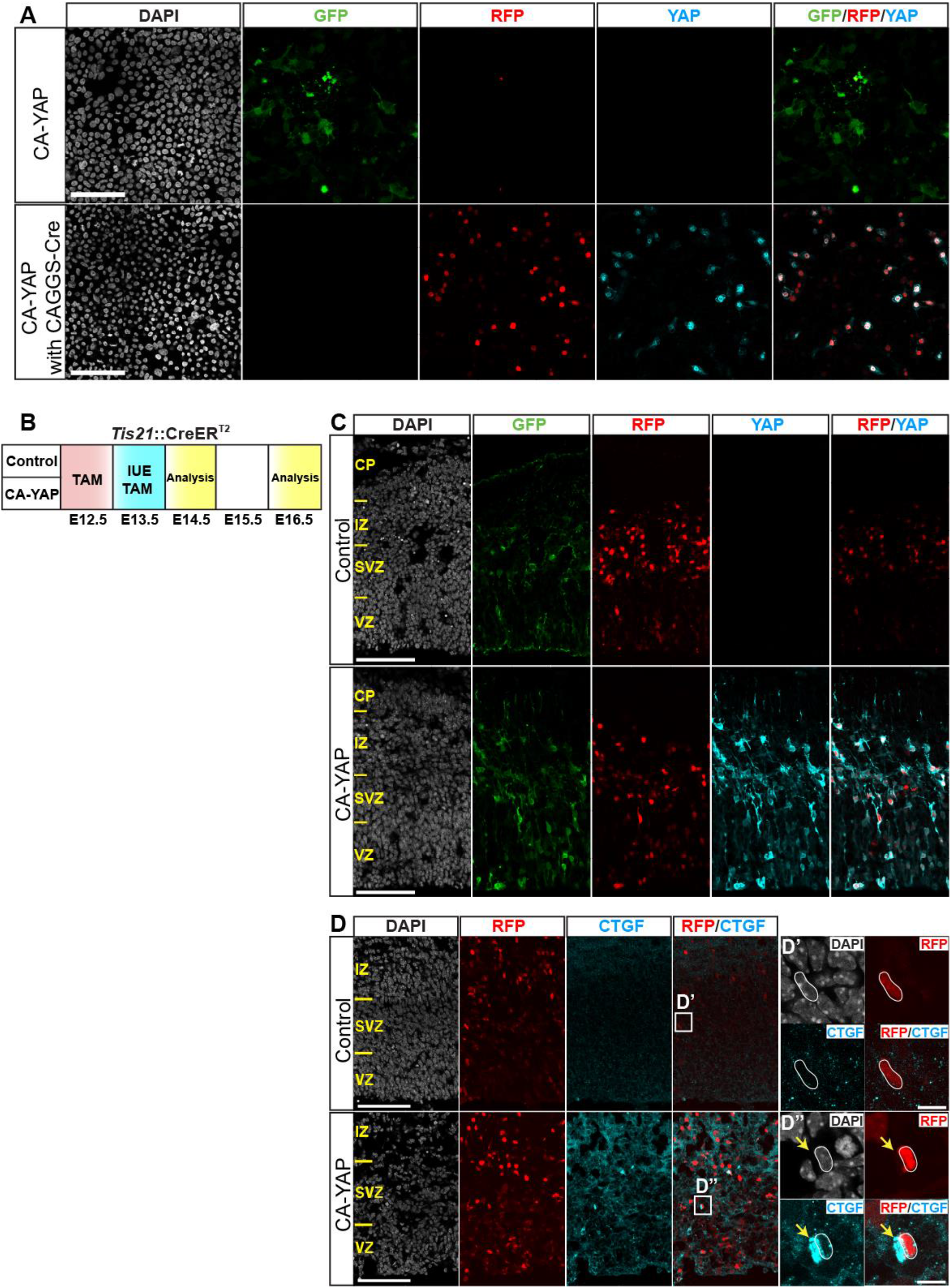
*In vitro* and *in vivo* validation of conditional CA-YAP expression. Related to Figures 2, 3 and 4. (A) *In vitro* validation. HEK293T cells were transfected with the CA-YAP plasmid (see Fig. 2A) only (top row), or with the CA-YAP plasmid together with a CAGGS-Cre plasmid (bottom row), followed by triple immunofluorescence for GFP (green), RFP (red) and YAP (cyan), combined with DAPI staining (white), 48 h later. (B) Flow scheme of *in vivo* validation experiment. *Tis21*::CreER^T2^ heterozygous mouse embryos received tamoxifen (TAM) at E12.5 and E13.5, and the neocortex was *in utero* electroporated (IUE) at E13.5 with control plasmid (C, D, top rows) or CA-YAP–expressing plasmid (C, D, bottom rows) (see Fig. 2A), followed by analysis at either E14.5 (C) or E16.5 (D). (C) Triple immunofluorescence for GFP (green), RFP (red) and YAP (cyan), combined with DAPI staining (white). (D) Double immunofluorescence for RFP (red) and CTGF (cyan), combined with DAPI staining (white). Boxes indicate areas in the SVZ that are shown at higher magnification in panels D’ and D”; white lines outline an RFP-positive nucleus in the control (D’) or upon conditional CA-YAP expression (D”); note the increase in CTGF immunoreactivity in the cytoplasm and adjacent extracellular space (arrows) upon conditional CA-YAP expression. (A, C, D) Images are 1-µm optical sections. Scale bars, 100 µm in (A, C, D), 10 µm in (D’, D”).

**Figure S4.**
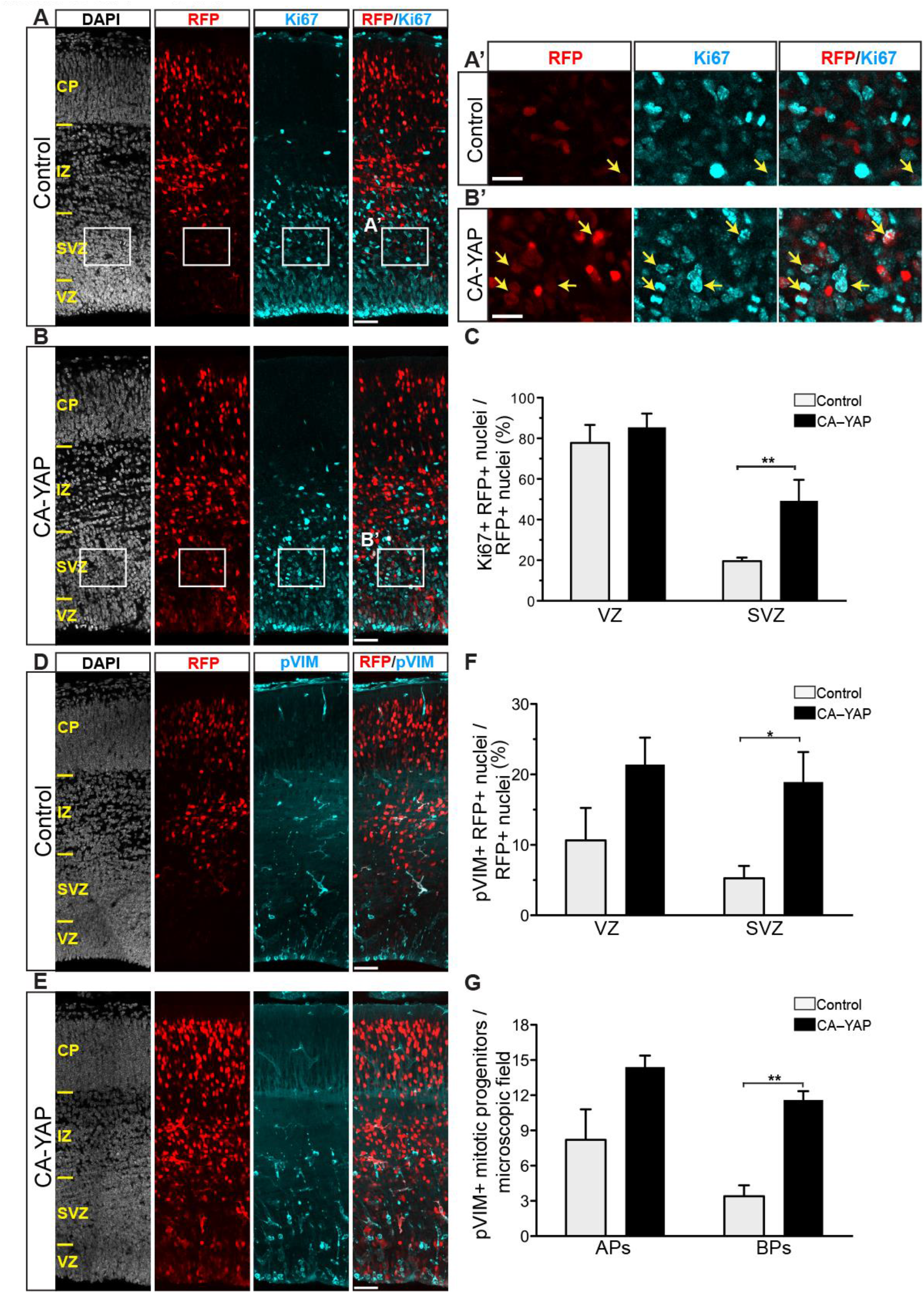
Conditional CA-YAP expression in the BP-genic lineage of embryonic mouse neocortex promotes BP proliferation. Related to Figure 3. *Tis21*::CreER^T2^ heterozygous mouse embryos received tamoxifen at E12.5 and E13.5, and the neocortex was *in utero* electroporated at E13.5 with control plasmid (A, A’, C, D, F, G) or CAYAP–expressing plasmid (B, B’ C, E, F, G) (see Fig. 2A), followed by analysis at E16.5. (A, B, D, E) Double immunofluorescence for RFP (red) and either Ki67 (A, B) or pVIM (D, E) (cyan), combined with DAPI staining (white). Boxes in (A, B) indicate areas in the SVZ that are shown at higher magnification in panels A’ and B’; arrows indicate selected RFP-positive nuclei that are Ki67-positive. Images are 1-µm optical sections. Scale bars, 50 µm in (A, B, D, E), 20 µm in (A’, B’). (C, G) Quantification of the percentage of RFP-positive cells that are Ki67-positive (C) and pVIM-positive (G) in the VZ and SVZ, upon control (light grey) and CA-YAP (black) electroporation. Two images (1-µm optical sections), each of 200 µm-wide field of cortical wall, per embryo were taken, and the percentage values obtained were averaged for each embryo. (F) Quantification of the number of APs and BPs in mitosis, as revealed by pVIM immunofluorescence, per microscopic field (200 µm-wide field of cortical wall), upon control (light grey) and CA-YAP (black) electroporation. Two images (1-µm optical sections) per embryo were taken, and the values obtained were averaged for each embryo. (C, F, G) Data are the mean of 5 embryos from five separate litters. Error bars indicate SEM; * *P* <0.05, ** *P* <0.01 (Mann-Whitney *U*-test).

**Figure S5.**
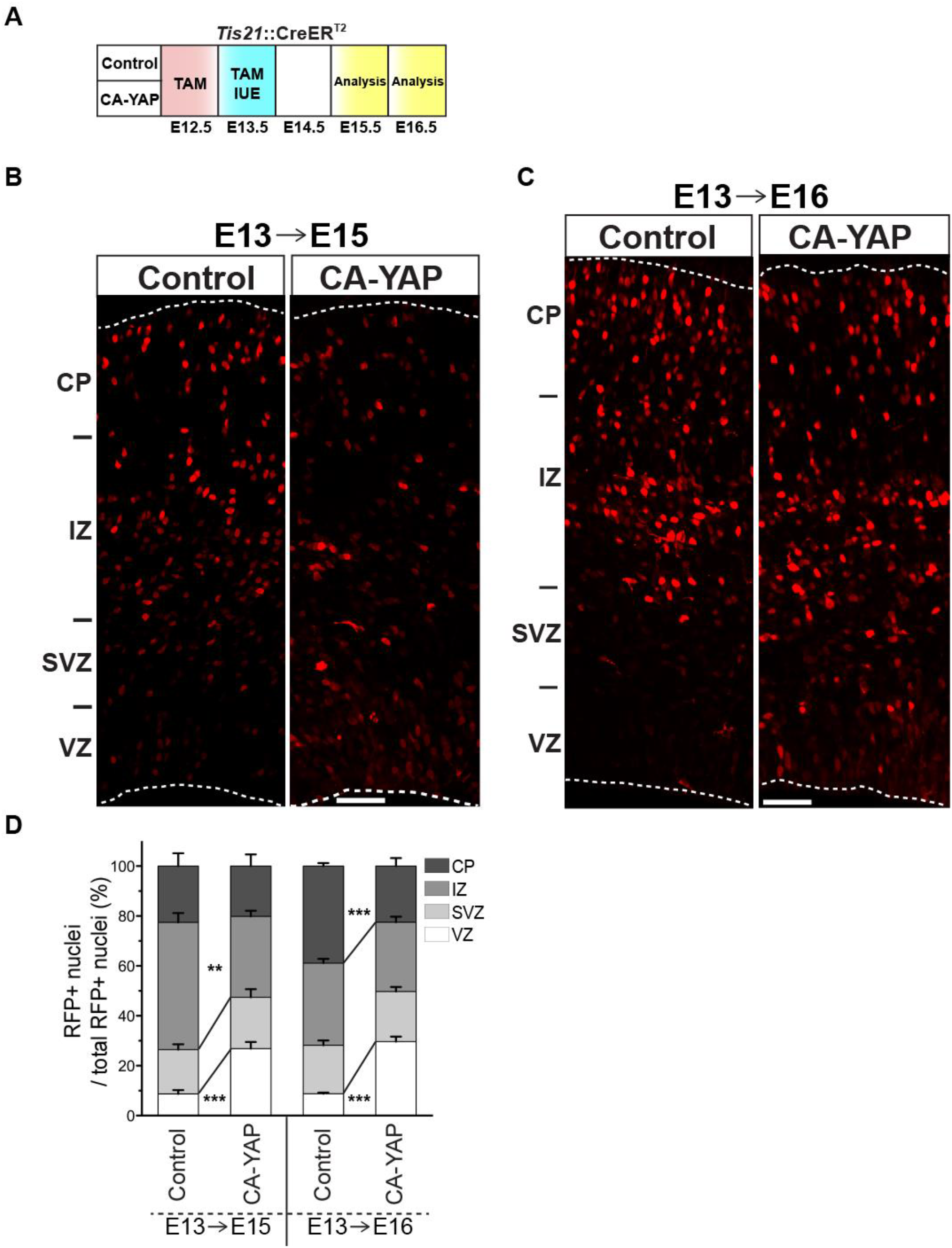
Conditional CA-YAP expression in the BP-genic lineage of embryonic mouse neocortex increases the relative abundance of progeny in the VZ. Related to Figures 2 and 3. (A) Flow scheme of experiments. *Tis21*::CreER^T2^ heterozygous mouse embryos received tamoxifen (TAM) at E12.5 and E13.5, and the neocortex was *in utero* electroporated (IUE) at E13.5 with control plasmid (B, C, left) or CA-YAP–expressing plasmid (B, C, right) (see Fig. 2A), followed by analysis at either E15.5 (B, D left) or E16.5 (C, D right). (B, C) Immunofluorescence for RFP (red) at either E15.5 (B) or E16.5 (C). Images are 1-µm optical sections. Scale bars, 50 µm. (D) Quantification of the percentage of total RFP-positive nuclei in the cortical wall that are found in the VZ, SVZ, IZ and CP, either 2 days (left two columns) or 3 days (right two columns) after control or CA-YAP electroporation. Two images (1-µm optical sections) per embryo were taken, each of a 200 µm-wide field of cortical wall, and the percentage values obtained were averaged for each embryo. Data are the mean of 5 embryos from five separate litters. Error bars indicate SEM; ** *P* <0.01, *** *P* <0.001 (unpaired Student’s *t-*test).

**Figure S6.**
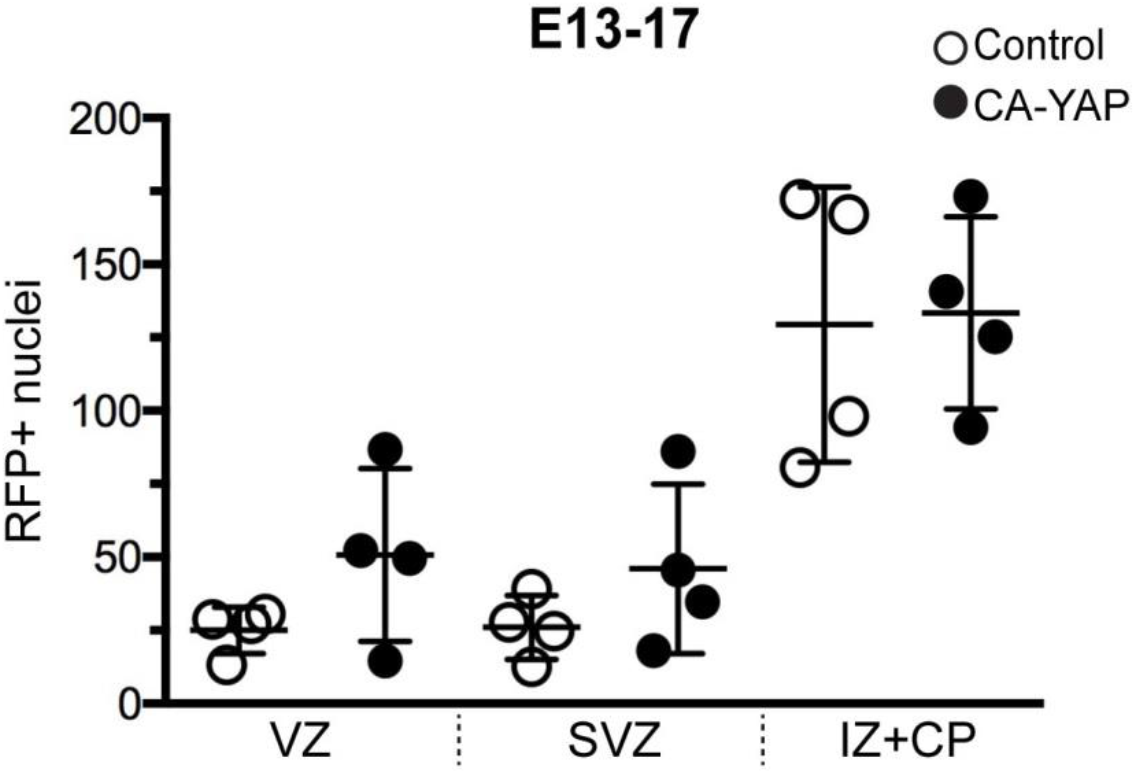
Number of RFP+ nuclei, four days upon IUE (E13-E17) in VZ, SVZ and IZ+CP. Related to Figure 4. Two images (1-µm optical sections), each of a 200 µm-wide field of cortical wall, per embryo were taken, and the percentage values obtained were averaged for each embryo. Data are the mean of 4 embryos from four separate litters.

### 2) Supplemental Results

#### *In vitro and in vivo* testing of CA-YAP plasmid conditional expression. Related to Figure S3

Conditional expression of CA-YAP was validated using two approaches, *in vitro* and *in vivo*. In the first, HEK293T were transfected with either CA-YAP–expressing plasmid (see Figure 2A) alone or CA-YAP– and CAGGS-Cre–expressing plasmids (Figure S3A). Whereas only membrane EGFP and no YAP expression was detected in CA-YAP–expressing plasmid alone (Figure S3A, upper row; note that fluorescent laser intensity is too low to detect endogenous YAP protein expression), upon addition of CAGGS-Cre–expressing plasmid, YAP and RFP reporter expressions were visible, with mainly nuclear YAP immunofluorescence (Figure S3A, lower row), suggesting an accurate recombination.

Further, to test *in vivo* expression, recombination and localization of CA-YAP, we electroporated embryonic neocortex of *Tis21*-CreER^T2+/-^ mouse line with CA-YAP–expressing plasmid (Figure S3B, C). The electroporation was performed at E13.5 and brains were analyzed one day later (Figure S3B). In the control, expression of CA-YAP was not detected (Figure S3C, upper row; note that fluorescent laser intensity is too low to detect endogenous YAP protein expression), whereas upon CA-YAP expression, YAP and RFP marker were mostly located in the VZ and SVZ (Figure S3C, lower row). This data shows successful recombination of CA-YAP–expressing plasmid *in* vivo and efficient delivery of CA-YAP to BPs of mouse embryonic neocortex.

In order to check if CA-YAP exerts its function, we analyzed expression of matricellular protein CTGF, a known transcriptional target of YAP (Malik et al., 2015; Zanconato et al., 2015; Zhao et al., 2008). Three days after conditional CA-YAP expression (Figure S3B), embryonic brain sections were immunostained for CTGF specific antibody. After CA-YAP expression, CTGF protein expression was found to be enriched and expressed in close proximity to RFP-positive nuclei in SVZ (Figure S3D, D”), whereas in control CTGF exhibited no expression in SVZ (Figure S3D, D’), as expected because also mRNA of *Ctgf* was absent in the SVZ of embryonic mouse E14.5 neocortex (Florio et al., 2015). This suggests that CA-YAP promoted expression of CTGF in the embryonic mouse SVZ. Taken together, these results show that this strategy is efficient for the conditional expression of functional CA-YAP in the neurogenic BPs in the SVZ.

### 3) Supplemental Experimental Procedures

#### YAP DNA constructs

Mouse wtYAP isoform 1 cDNA clone (IMAGE 4239820) was obtained from Source Bioscience. Restriction enzyme sites, SalI, were adapted to the 5’ and 3’ ends of wtYAP by PCR. wtYAP was digested by SalI and subcloned to the *pCR2.1-TOPO-TA* vector (Invitrogen), creating an intermediate, non-expressing vector *pTOPO-mouse-wtYAP*. The destination vector *pCAGGSLoxP-Gap43-GFP-LoxP-IRES-nRFP* (Wong et al., 2015) was used as control plasmid and used to obtain the CA-YAP plasmid. To this end, it was digested by XhoI, and *pTOPO-mouse-wtYAP* was digested by SalI to obtain linearized wtYAP DNA. The opened destination vector (*pCAGGS–LoxP–Gap43-GFP–LoxP–IRES–nRFP*) and the linearized wtYAP DNA were purified by 2% agarose gel electrophoresis and ligated together to yield *pCAGGS–LoxP–Gap43-GFP– LoxP–wtYAP–IRES–nRFP*. This plasmid was then used to generate the CA-YAP-carrying plasmid (*pCAGGS–LoxP–Gap43-GFP–LoxP–CA-YAP–IRES–nRFP*) by replacing two serine residues, YAP-S112 and YAP-S382, with alanine residues, by PCR point mutagenesis.

To obtain the DN-YAP plasmid, *pcDNA3.1-pA83-dnYAP* (RBD12195) was obtained from the RIKEN BRC through the National Bio-Resource Project of the MEXT, Japan. In *pcDNA3.1-pA83-dnYAP*, YAP-S112 is replaced with alanine, and the transactivation domain of mouse YAP is replaced with the engrailed repression domain from *Drosophila*. *pcDNA3.1-pA83-dnYAP* was subcloned into the *pCR2.1-TOPO-TA* vector (Invitrogen) using PCR and adding BamHI sites to the 5’ and 3’ ends of dnYAP, creating an intermediate, non-expressing vector *pTOPO-mouse-DN-YAP*. The destination vector *pCAGGS-empty* (Niwa et al., 1991) was used as control plasmid and used to obtain the DN-YAP plasmid. To this end, it was digested by BglII, and *pTOPO-mouse-DN-YAP* was digested by BamHI to obtain linearized DN-YAP DNA. The opened destination vector (*pCAGGS-empty*) and the linearized DN-YAP DNA were purified by 2% agarose gel electrophoresis and ligated together to yield *pCAGGS-DN-YAP*.

Primers for cloning mouse YAP:

YAP-SalI-F: 5’-gcgcgtcgacgccaccatggagcccgcgcaacagcc-3’

YAP-SalI-R: 5’-cgcggtcgacctataaccacgtgagaaagct-3’

Primers for generating point mutations:

S112 → A112

S112A-F: 5’-catgttcgagctcacgcctctccagcctccc-3’

S112A-R: 5’-gggaggctggagaggcgtgagctcgaacatg-3’

S382 → A382

S382A-F: 5’-cactctcgagatgaggccacagacagcggcc-3’

S382A-R: 5’-ggccgctgtctgtggcctcatctcgagagtg-3’

Primers for cloning DN-YAP:

BamHI-HA-F: 5’-gcgcggatccgccaccatgtacccatacgacgttcc-3’

BamHI-HA-R: 5’-cgcgggatccctagttcaggtcctcctcgg-3’

#### HEK293T cell transfection

HEK293T cells were grown in DMEM (Gibco) supplemented with 10% fetal calf serum and containing 1% penicillin/streptomycin (Gibco 15140122) at 37°C in an atmosphere of 5% CO_2_ / 95% air. The transfection was performed with Lipofectamine 2000 reagent (Invitrogen). The day before transfection, cells were plated on a 24-well plate, 10^5^ cells per well, in the above cell culture medium. Cells were either lipofectamine transfected with CA-YAP only (600 ng per well) or co-transfected with CA-YAP and CAGGS-Cre (each 600 ng per well). After 48 h, cells were harvested and fixed in 4% PFA for 20 min. The cells were washed with 1x PBS and processed for further analysis within 24 h.

#### EdU labeling

To analyze cell-cycle re-entry, one pulse of EdU was administered by intraperitoneal injection of 0.1 ml of EdU (1 mg/ml) into pregnant mice carrying E14.5 embryos, 24 h before the animal was sacrificed. At this stage of mouse cortical neurogenesis, the average length of the sum of S-phase, G2 plus M-phase of *Tis21*-GFP–positive APs and BPs is ≈4 h and ≈5 h, respectively (Arai et al., 2011). Hence, the time period of 24 h between the EdU pulse and the analysis is sufficient for the incorporated EdU to become inherited by daughter cells. To determine whether or not an EdU-labeled daughter-cell (see below for EdU detection) derived from an RFP-expressing mother cell re-entered the cell-cycle, Ki67 immunofluorescence was performed as is described below. EdU detection was performed on cryosections. After the incubation with secondary antibodies (see below), the tissue was fixed again, for 20 min with 4% PFA. EdU detection was carried out following the protocol of the Click-iT EdU kit with Alexa Fluor 647 (Invitrogen), as described previously (Arai et al., 2011).

#### Immunofluorescence on fixed cells and tissues

Transfected HEK293T cells were incubated with trypsin (Gibco 25300054) (5 min), harvested, sedimented (300 x *g*), resuspended in 1x PBS, and fixed in 4% PFA for 10 min at room temperature. Cells were permeabilized with 0.3% Triton X-100 in 1x PBS for 10 min, followed by quenching in 0.1 M glycine in 1x PBS for 10 min. Primary antibodies were incubated for 2 h at room temperature, followed by incubation with secondary antibodies for 1 h, all in 1x PBS containing 0.2% gelatin, an additional 300 mM NaCl, and 0.3% Triton X-100 (PGNT buffer). Coverslips with the fixed, permeabilized and immunostained cells were washed with 1x PBS and mounted on glass slides using Mowiol.

Embryonic mouse and ferret brain and fetal human brain tissues fixed in 4% PFA were washed in 1x PBS, immersed in 30% sucrose, and kept overnight in sucrose at 4°C on a rocking platform. The sucrose-soaked tissue was embedded with Tissue-Tek solution (O.C.T., Sakura Finetek) and frozen at –20°C. Tissue was sectioned on a cryostat (20-µm cryosections). Frozen cryosections were rehydrated in 1x PBS. To be able to reliably detect nuclear epitopes, we routinely carried out an antigen retrieval protocol as follows. Cryosections were heated in 0.01 M Na-citrate pH 6.0 at 70°C for 45-60 min. After cooling to room temperature cryosections were permeabilized with 0.3% Triton X-100 in the 1x PBS for 30 min and quenched in 0.1 M glycine in 1x PBS for 30 min. Primary antibodies were incubated overnight at 4°C, followed by incubation with secondary antibodies for 2 h, all in PGNT buffer. After several washes in PGNT buffer and then in 1x PBS, cryosections were mounted on glass slides using Mowiol.

The following primary antibodies were used: CTGF (goat, polyclonal, Santa Cruz, 14939, 1:100), GFP (chicken, polyclonal, Aves, GFP1020, 1:500), Ki67 (rabbit, polyclonal, Abcam, ab15580, 1:100), PH3-S28 (rat, monoclonal, Abcam, ab10543, 1:250), pVIM (mouse, monoclonal, Abcam, ab22651, 1:100), RFP (rat, monoclonal, Chromotek, 5F8, 1:500), Satb2 (mouse, monoclonal, Abcam, ab51502, 1:100), Sox2 (goat, polyclonal, Santa Cruz, SC17320, 1:100), Sox2 (goat, polyclonal, R&D Systems, AF2018, 1:100), Tbr1 (rabbit, polyclonal, Abcam, ab31940, 1:100), Tbr2 (rabbit, polyclonal, Abcam, ab23345, 1:200), Tbr2 (mouse, MPI-CBG Antibody Facility, 1:50), phospho-YAP-S127 (rabbit, polyclonal, Cell Signaling, 4911, 1:50), phospho-YAP-S127 (rabbit, monoclonal, Cell Signaling, 13008, 1:100), YAP (rabbit, monoclonal, Cell Signaling, 14074, 1:100), and YAP (mouse, monoclonal, Abcam, ab56701, 1:100). Alexa Fluor 488, 555, 647 (donkey, Molecular Probes) and Cy2, Cy3, Cy5 (Jackson) labeled secondary antibodies were used (1:500). Nuclei were counterstained with DAPI (Sigma, 1:1000).

#### Image acquisition

All images were acquired using a Carl Zeiss LSM 700 confocal light microscope. Objectives 40X (oil) and 25X were used. Images were taken as 1-µm optical sections in Z-stack and tile scans which were post-stitched in ZEN software (Carl Zeiss). All images presented are 1-µm optical sections.

#### Determination of germinal zones, apical and basal mitoses

Germinal zones were determined according to the differences in the cytoarchitecture as revealed by DAPI staining of nuclei. The VZ was defined as the zone of a pseudostratified epithelium where nuclei are elongated, densely packed and radially aligned. The SVZ was defined as the zone of non-radially aligned, rounded and less densely packed nuclei. In ferret and human, iSVZ and oSVZ were distinguished according to the density of nuclei, where the iSVZ comprises densely packed rounded nuclei and the oSVZ comprises sparse nuclei. The IZ was defined as the zone located between SVZ and CP, which had sparser nuclei than the SVZ (or the oSVZ in ferret and human). The CP was defined as the zone of densely packed rounded nuclei beneath the pial surface. Apical mitoses were defined as pVIM- or PH3-positive mitoses occurring within the VZ, and basal mitoses were defined as pVIM- or PH3-positive mitoses occurring within the SVZ (Figure S4F,G; Figure 5B,D; Figure 6C,E,F).

#### Quantifications

All quantifications were performed on 1-µm optical sections, with the exception of pVIM staining in Figure 6E and F where 5 optical sections were analyzed (5-µm Z-stack). Quantifications were performed using the Fiji software with the cell counter plugin and/or “measure” function.

For Figure 2D,G; Figure 3D,H; Figure 4F,G; Figure S4D; and Figure S4C,F, nuclei were quantified based on the expression of the respective marker combined with RFP expression. The scored double-positive nuclei (RFP plus selected marker) were expressed as a percentage of the total number of RFP-positive nuclei in the given germinal zone. Quantification was done on 200-µm wide images, oriented parallel to the apical surface.

Quantifications of RFP-positive nuclei in each zone (Figure S5D) were expressed as a percentage of the number of RFP-positive nuclei in the given zone over the total number RFP-positive nuclei in the cortical wall.

To determine if a nucleus is YAP-positive (Figure 1), we measured and averaged the YAP immunofluorescence intensity for 10 nuclei located in the CP as background. In the VZ or SVZ, we scored a nucleus as YAP-positive if the immunofluorescence intensity was at least two times higher than the background immunofluorescence. For quantification of YAP immunoreactivity, we acquired two-three images per embryo/fetus, and in each image (i.e., each cryosection) 30 randomly selected DAPI-positive (Figure 1D), Sox2-positive (Figure 1E), Tbr2-negative or -positive (Figure 1I-K) nuclei in the VZ (Figure 1I,J) or in the SVZ (Figure 1D,E,K) were scored. The values obtained were averaged for each embryo/fetus. Data are expressed as the percentage of 30 scored cells.

#### Total YAP vs. phospho-YAP

For the comparison of total YAP vs. phospho-YAP immunoreactivity and the determination of nuclear dephospho-YAP levels (Figure S2A and B), immunofluorescence was performed as follows. Cryosections of fixed mouse, ferret and human neocortex of the indicated developmental stages (1-2 cryosections per embryo/fetus) were incubated overnight at 4°C with two primary antibodies against YAP, a mouse monoclonal antibody recognizing total YAP (Abcam, ab56701, 1:100) and a rabbit polyclonal antibody recognizing only YAP phosphorylated at serine127 (phospho-YAP, Cell Signaling, 4911, 1:100), and a polyclonal goat antibody against Sox2 (R&D Systems, AF2018, 1:100), followed by incubation with the respective appropriate secondary antibodies, anti-mouse-Alexa Fluor 488, anti-rabbit Alexa Fluor 555 and anti-goat Alexa Fluor 647 (Molecular Probes), for 2 h at room temperature. After a series of washes in 1x PBS, sections were mounted on glass slides using Mowiol.

For the quantification of YAP immunoreactivity, one image per cryosection was taken and 30 randomly selected Sox2-positive nuclei in the SVZ were scored per cryosection. For both, the total YAP channel and the phospho-YAP channel, immunofluorescence background values were determined by averaging the fluorescence signals of 10 DAPI-stained nuclei in the CP per image (as CP nuclei lacked YAP immunoreactivity, see Figure 1); these background values were then subtracted from the respective total YAP and phospho-YAP immunofluorescence values obtained for each of the Sox2-positive nuclei per cryosection. To be able to relate the resulting, background-corrected, nuclear total YAP and nuclear phospho-YAP immunofluorescence values to each other, the mean immunofluorescence values for total YAP and for phospho-YAP from three representative areas of cytoplasm per image were determined. The ratio of mean cytoplasmic total YAP immunofluorescene value / mean cytoplasmic phospho-YAP immunofluorescene value was multiplied with the nuclear phospho-YAP immunofluorescence values to yield the adjusted immunofluorescence values for nuclear phospho-YAP. Then, for each nucleus, the adjusted immunofluorescence value for phospho-YAP was subtracted from the_immunofluorescence value for total YAP, to yield the value for dephosphorylated, i.e. active, YAP.

#### Protein phosphatase treatment

To determine the proportion of YAP in Sox2+ nuclei in the SVZ that was in dephosphorylated form and hence active (Figure S2C), immunofluorescence was performed as follows. Cryosections (3 µm) of fixed mouse, ferret and human neocortex of the indicated developmental stages (1-2 cryosections per embryo/fetus) were subjected to antigen retrieval followed by permeabilization as described above. Cryosections were then treated with lambda protein phosphatase (New England Biolabs, P0753S) in a total reaction volume of 100 µl per cryosection, which consisted of 10 µl of 10x Protein MetalloPhosphatase buffer (50 mM HEPES pH 7.5, 10 mM NaCl, 2 mM DTT, 0.01% Brij 35; New England Biolabs, B0761S), 10 µl of 10 mM MnCl_2_, and either 10 µl of 400 units/µl of lambda protein phosphatase plus 70 µl of water (phosphatase treatment), or 80 µl of water (control treatment). For each neocortex sample per species, three cryosections were subjected to phosphatase treatment for 2 h at 37°C in a humidified chamber, and three other cryosections were subjected to control treatment. Cryosections were then washed in 0.3% Triton X-100 in 1x Tris-HCl buffered saline containing 0.2% gelatine (washing/blocking buffer), and blocked for 30 min in the same buffer followed by overnight incubation at 4°C with a mixture of two primary rabbit monoclonal antibodies diluted in washing/blocking buffer, one to detect total YAP (Cell Signaling, 14074, 1:100) and the other to detect phospho-YAP (Cell Signaling, 13008, 1:100), and polyclonal goat Sox2 antibody (R&D Systems, AF2018, 1:100), followed by incubation in washing/blocking buffer containing the respective appropriate secondary antibodies, anti-rabbit Alexa Fluor 488 and anti-goat Alexa Fluor 555 (Molecular Probes), for 2 h at room temperature. After a series of washes in 1x PBS, cryosections were mounted to glass slides using Mowiol.

For the quantification of the YAP immunoreactivity detected with the mixture of the total YAP plus phospho-YAP antibodies upon control vs. phosphatase treatment, one image per cryosection was taken and 30 randomly selected Sox2-positive nuclei in the SVZ were scored per cryosection. For each neocortex sample per species, after determining the average value per cryosection, the mean of these average values for the three control cryosections was set to 100%, and the mean of the average values for the three phosphatase-treated cryosections was expressed relative to this. The reduction, upon protein phosphatase treatment, in the YAP immunofluorescence signal obtained with the sum of the two antibodies (total YAP plus phospho-YAP) indicates the contribution of phospho-YAP to this signal.

## References

Arai, Y., Pulvers, J.N., Haffner, C., Schilling, B., Nusslein, I., Calegari, F., and Huttner, W.B. (2011). Neural stem and progenitor cells shorten S-phase on commitment to neuron production. Nat. Commun. 2, 154.

Attardo, A., Calegari, F., Haubensak, W., Wilsch-Brauninger, M., and Huttner, W.B. (2008). Live imaging at the onset of cortical neurogenesis reveals differential appearance of the neuronal phenotype in apical versus basal progenitor progeny. PLoS One 3, e2388.

Barry, E.R., and Camargo, F.D. (2013). The Hippo superhighway: signaling crossroads converging on the Hippo/Yap pathway in stem cells and development. Curr. Opin. Cell. Biol. 25, 247-253.

Basu-Roy, U., Bayin, N.S., Rattanakorn, K., Han, E., Placantonakis, D.G., Mansukhani, A., and Basilico, C. (2015). Sox2 antagonizes the Hippo pathway to maintain stemness in cancer cells. Nat. Commun. 6, 6411-6429.

Brodowska, K., Al-Moujahed, A., Marmalidou, A., Meyer Zu Horste, M., Cichy, J., Miller, J.W., Gragoudas, E., and Vavvas, D.G. (2014). The clinically used photosensitizer Verteporfin (VP) inhibits YAP-TEAD and human retinoblastoma cell growth in vitro without light activation. Exp. Eye Res. 124, 67-73.

Camargo, F.D., Gokhale, S., Johnnidis, J.B., Fu, D., Bell, G.W., Jaenisch, R., and Brummelkamp, T.R. (2007). YAP1 increases organ size and expands undifferentiated progenitor cells. Curr. Biol. 17, 2054-2060.

Cappello, S., Gray, M.J., Badouel, C., Lange, S., Einsiedler, M., Srour, M., Chitayat, D., Hamdan, F.F., Jenkins, Z.A., Morgan, T., et al. (2013). Mutations in genes encoding the cadherin receptor-ligand pair DCHS1 and FAT4 disrupt cerebral cortical development. Nat. Genet. 45, 1300-1308.

De Juan Romero, C., Bruder, C., Tomasello, U., Sanz-Anquela, J.M., and Borrell, V. (2015). Discrete domains of gene expression in germinal layers distinguish the development of gyrencephaly. EMBO J. 34, 1859-1874.

Dehay, C., Kennedy, H., and Kosik, K.S. (2015). The outer subventricular zone and primate-specific cortical complexification. Neuron 85, 683-694.

Dong, J., Feldmann, G., Huang, J., Wu, S., Zhang, N., Comerford, S.A., Gayyed, M.F., Anders, R.A., Maitra, A., and Pan, D. (2007). Elucidation of a universal size-control mechanism in Drosophila and mammals. Cell 130, 1120-1133.

Englund, C., Fink, A., Lau, C., Pham, D., Daza, R.A., Bulfone, A., Kowalczyk, T., and Hevner, R.F. (2005). Pax6, Tbr2, and Tbr1 are expressed sequentially by radial glia, intermediate progenitor cells, and postmitotic neurons in developing neocortex. J. Neurosci. 25, 247-251.

Fietz, S.A., and Huttner, W.B. (2011). Cortical progenitor expansion, self-renewal and neurogenesis-a polarized perspective. Curr. Opin. Neurobiol. 21, 23-35.

Fietz, S.A., Kelava, I., Vogt, J., Wilsch-Brauninger, M., Stenzel, D., Fish, J.L., Corbeil, D., Riehn, A., Distler, W., Nitsch, R., et al. (2010). OSVZ progenitors of human and ferret neocortex are epithelial-like and expand by integrin signaling. Nat. Neurosci. 13, 690-699.

Fietz, S.A., Lachmann, R., Brandl, H., Kircher, M., Samusik, N., Schroder, R., Lakshmanaperumal, N., Henry, I., Vogt, J., Riehn, A., et al. (2012). Transcriptomes of germinal zones of human and mouse fetal neocortex suggest a role of extracellular matrix in progenitor self-renewal. Proc. Natl. Acad. Sci. USA 109, 11836-11841.

Florio, M., Albert, M., Taverna, E., Namba, T., Brandl, H., Lewitus, E., Haffner, C., Sykes, A., Wong, F.K., Peters, J., et al. (2015). Human-specific gene ARHGAP11B promotes basal progenitor amplification and neocortex expansion. Science 347, 1465-1470.

Florio, M., and Huttner, W.B. (2014). Neural progenitors, neurogenesis and the evolution of the neocortex. Development 141, 2182-2194.

Geschwind, D.H., and Rakic, P. (2013). Cortical evolution: judge the brain by its cover. Neuron 80, 633-647.

Hansen, D.V., Lui, J.H., Parker, P.R., and Kriegstein, A.R. (2010). Neurogenic radial glia in the outer subventricular zone of human neocortex. Nature 464, 554-561.

Hao, Y., Chun, A., Cheung, K., Rashidi, B., and Yang, X. (2007). Tumor suppressor LATS1 Is a negative regulator of oncogene YAP. J. Biol. Chem. 283, 5496-5509.

Haubensak, W., Attardo, A., Denk, W., and Huttner, W.B. (2004). Neurons arise in the basal neuroepithelium of the early mammalian telencephalon: a major site of neurogenesis. Proc. Natl. Acad. Sci. USA 101, 3196-3201.

Kawasaki, H., Iwai, L., and Tanno, K. (2012). Rapid and efficient genetic manipulation of gyrencephalic carnivores using in utero electroporation. Mol. Brain 5, 24.

Kawasaki, H., Toda, T., and Tanno, K. (2013). In vivo genetic manipulation of cortical progenitors in gyrencephalic carnivores using in utero electroporation. Biol. Open 2, 95-100.

Kelava, I., Reillo, I., Murayama, A.Y., Kalinka, A.T., Stenzel, D., Tomancak, P., Matsuzaki, F., Lebrand, C., Sasaki, E., Schwamborn, J.C., et al. (2012). Abundant occurrence of basal radial glia in the subventricular zone of embryonic neocortex of a lissencephalic primate, the common marmoset Callithrix jacchus. Cereb. Cortex 22, 469-481.

Kowalczyk, T., Pontious, A., Englund, C., Daza, R.A., Bedogni, F., Hodge, R., Attardo, A., Bell, C., Huttner, W.B., and Hevner, R.F. (2009). Intermediate neuronal progenitors (basal progenitors) produce pyramidal-projection neurons for all layers of cerebral cortex. Cereb. Cortex 19, 2439-2450.

Lavado, A., He, Y., Pare, J., Neale, G., Olson, E.N., Giovannini, M., and Cao, X. (2013). Tumor suppressor Nf2 limits expansion of the neural progenitor pool by inhibiting Yap/Taz transcriptional coactivators. Development 140, 3323-3334.

Lavado, A., Ware, M., Paré, J., and Cao, X. (2014). The tumor suppressor Nf2 regulates corpus callosum development by inhibiting the transcriptional coactivator Yap. Development 141, 4182-4193.

Lian, I., Kim, J., Okazawa, H., Zhao, J., Zhao, B., Yu, J., Chinnaiyan, A., Israel, M.A., Goldstein, L.S., Abujarour, R., et al. (2010). The role of YAP transcription coactivator in regulating stem cell self-renewal and differentiation. Genes Dev. 24, 1106-1118.

Liu-Chittenden, Y., Huang, B., Shim, J.S., Chen, Q., Lee, S.J., Anders, R.A., Liu, J.O., and Pan, D. (2012). Genetic and pharmacological disruption of the TEAD-YAP complex suppresses the oncogenic activity of YAP. Genes Dev. 26, 1300-1305.

Long, K.R., Newland, B., Florio, M., Kalebic, N., Langen, B., Kolterer, A., Wimberger, P., and Huttner, W.B. (2018). Extracellular matrix components HAPLN1, lumican and collagen I cause hyaluronic acid-dependent folding of the developing human neocortex. Neuron 99, 702-719.

Lui, J.H., Hansen, D.V., and Kriegstein, A.R. (2011). Development and evolution of the human neocortex. Cell 146, 18-36.

Malik, A.R., Liszewska, E., and Jaworski, J. (2015). Matricellular proteins of the Cyr61/CTGF/NOV (CCN) family and the nervous system. Front. Cell Neurosci. 9, 237.

Miyata, T., Kawaguchi, A., Saito, K., Kawano, M., Muto, T., and Ogawa, M. (2004). Asymmetric production of surface-dividing and non-surface-dividing cortical progenitor cells. Development 131, 3133-3145.

Namba, T., and Huttner, W.B. (2017). Neural progenitor cells and their role in the development and evolutionary expansion of the neocortex. WIREs Dev. Biol. 6, e256.

Nishioka, N., Inoue, K., Adachi, K., Kiyonari, H., Ota, M., Ralston, A., Yabuta, N., Hirahara, S., Stephenson, R.O., Ogonuki, N., et al. (2009). The Hippo signaling pathway components Lats and Yap pattern Tead4 activity to distinguish mouse trophectoderm from inner cell mass. Dev.Cell 16, 398-410.

Noctor, S.C., Martinez-Cerdeno, V., Ivic, L., and Kriegstein, A.R. (2004). Cortical neurons arise in symmetric and asymmetric division zones and migrate through specific phases. Nat. Neurosci. 7, 136-144.

Pontious, A., Kowalczyk, T., Englund, C., and Hevner, R.F. (2008). Role of intermediate progenitor cells in cerebral cortex development. Dev. Neurosci. 30, 24-32.

Rakic, P. (2009). Evolution of the neocortex: a perspective from developmental biology. Nat. Rev. Neurosci. 10, 724-735.

Reillo, I., de Juan Romero, C., Garcia-Cabezas, M.A., and Borrell, V. (2011). A role for intermediate radial glia in the tangential expansion of the mammalian cerebral cortex. Cereb. Cortex 21, 1674-1694.

Saito, K., Kawasoe, R., Sasaki, H., Kawaguchi, A., and Miyata, T. (2017). Neural progenitor cells undergoing Yap/Tead-Mediated enhanced self-renewal form heterotopias more easily in the diencephalon than in the telencephalon. Neurochem. Res. 43, 180-189.

Schenk, J., Wilsch-Brauninger, M., Calegari, F., and Huttner, W.B. (2009). Myosin II is required for interkinetic nuclear migration of neural progenitors. Proc. Natl. Acad. Sci. USA 106, 16487-16492.

Seo, E., Basu-Roy, U., Gunaratne, P.H., Coarfa, C., Lim, D.S., Basilico, C., and Mansukhani, A. (2013). SOX2 regulates YAP1 to maintain stemness and determine cell fate in the osteoadipo lineage. Cell Rep. 3, 2075-2087.

Shitamukai, A., Konno, D., and Matsuzaki, F. (2011). Oblique radial glial divisions in the developing mouse neocortex induce self-renewing progenitors outside the germinal zone that resemble primate outer subventricular zone progenitors. J. Neurosci. 31, 3683-3695.

Smart, I.H., Dehay, C., Giroud, P., Berland, M., and Kennedy, H. (2002). Unique morphological features of the proliferative zones and postmitotic compartments of the neural epithelium giving rise to striate and extrastriate cortex in the monkey. Cereb. Cortex 12, 37-53.

Song, S., Ajani, J.A., Honjo, S., Maru, D.M., Chen, Q., Scott, A.W., Heallen, T.R., Xiao, L., Hofstetter, W.L., Weston, B., et al. (2014). Hippo coactivator YAP1 upregulates SOX9 and endows esophageal cancer cells with stem-like properties. Cancer Res. 74, 4170-4182.

Sudol, M., Shields, D.C., and Farooq, A. (2012). Structures of YAP protein domains reveal promising targets for development of new cancer drugs. Semin. Cell Dev. Biol. 23, 827-833.

Turrero Garcia, M., Chang, Y., Arai, Y., and Huttner, W.B. (2015). S-phase duration is the main target of cell cycle regulation in neural progenitors of developing ferret neocortex. J. Comp. Neurol. 524, 456-470.

Wang, C., Zhu, X., Feng, W., Yu, Y., Jeong, K., Guo, W., Lu, Y., and Mills, G.B. (2016). Verteporfin inhibits YAP function through up-regulating 14-3-3sigma sequestering YAP in the cytoplasm. AM. J. Cancer Res. 6, 27-37.

Wang, X., Tsai, J.W., LaMonica, B., and Kriegstein, A.R. (2011). A new subtype of progenitor cell in the mouse embryonic neocortex. Nat. Neurosci. 14, 555-561.

Wilsch-Bräuninger, M., Florio, M., and Huttner, W.B. (2016). Neocortex expansion in development and evolution - from cell biology to single genes. Curr. Opin. Neurobiol. 39, 122-132.

Wong, F.K., Fei, J.F., Mora-Bermudez, F., Taverna, E., Haffner, C., Fu, J., Anastassiadis, K., Stewart, A.F., and Huttner, W.B. (2015). Sustained Pax6 expression generates primate-like basal radial glia in developing mouse neocortex. PLoS Biol. 13, e1002217.

Yu, F.X., Zhao, B., and Guan, K.L. (2015). Hippo pathway in organ size control, tissue homeostasis, and cancer. Cell 163, 811-828.

Zanconato, F., Forcato, M., Battilana, G., Azzolin, L., Quaranta, E., Bodega, B., Rosato, A., Bicciato, S., Cordenonsi, M., and Piccolo, S. (2015). Genome-wide association between YAP/TAZ/TEAD and AP-1 at enhancers drives oncogenic growth. Nat. Cell Biol. 17, 1218-1227.

Zhao, B., Li, L., Tumaneng, K., Wang, C.Y., and Guan, K.L. (2010). A coordinated phosphorylation by Lats and CK1 regulates YAP stability through SCF(beta-TRCP). Genes Dev. 24, 72-85.

Zhao, B., Wei, X., Li, W., Udan, R.S., Yang, Q., Kim, J., Xie, J., Ikenoue, T., Yu, J., Li, L., et al. (2007). Inactivation of YAP oncoprotein by the Hippo pathway is involved in cell contact inhibition and tissue growth control. Genes Dev. 21, 2747-2761.

Zhao, B., Ye, X., Yu, J., Li, L., Li, W., Li, S., Yu, J., Lin, J.D., Wang, C.Y., Chinnaiyan, A.M., et al. (2008). TEAD mediates YAP-dependent gene induction and growth control. Genes Dev. 22, 1962-1971.

## Supplemental References

Niwa, H., Yamamura, K., and Miyazaki, J. (1991). Efficient selection for high-expression transfectants with a novel eukaryotic vector. Gene 108, 193-199.

